# Systemic iron sequestration restricts the intratumoral CD8^+^ T cell landscape in pancreas cancer

**DOI:** 10.64898/2026.09.21.753167

**Authors:** Joshua D. Schoenfeld, Jahi J. Noel, Christopher S. McGinnis, Chenlian Fu, Brinda Alagesan, Eduardo Hernandez, Justin Jee, Nikolaus Schultz, Quaid Morris, Scott Lowe, Suzanne M. Cloonan, John E. Connolly, Ronan Chaligne, David Kelsen, Kenneth H. Yu, William Z. Zhang, Eileen M. O’Reilly, Ansuman T. Satpathy, Tomas Ganz, Santosha A. Vardhana

## Abstract

Systemic iron sequestration occurs frequently in cancer due to inflammation-driven expression of the iron-regulatory hormone hepcidin. The impact of systemic iron availability on tumoral immunity is unclear. Here, we show that elevated serum hepcidin is associated with reduced survival and decreased intratumoral CD8^+^ T cells in patients with pancreas cancer. While hepcidin is not induced in murine tumor models, administration of a hepcidin mimetic phenocopies the T-cell-depleted tumor microenvironment seen in patients. Mechanistically, chronic antigen-driven mitochondrial dysfunction disrupts iron metabolism and selectively depletes high avidity CD8^+^ T cells during iron restriction. These findings establish a direct link between hepcidin-mediated iron sequestration and tumoral immunity and nominate systemic iron dysregulation as a therapeutic target to enhance anti-tumoral CD8^+^ T cell responses.

## Main Text

Disruptions of systemic iron metabolism in patients with cancer were first described in the 1970s when serum ferritin was proposed as a diagnostic biomarker (*1, 2*). It is now understood that increased serum ferritin reflects the sequestration of systemic iron in both cancer and non-malignant inflammatory conditions and may be associated with functional iron deficiency and anemia of chronic disease (*3*). In contrast to absolute iron deficiency, which reflects decreased total body iron stores, functional iron deficiency occurs primarily in response to hepcidin, an acute phase reactant that hormonally restricts systemic iron availability in response to inflammatory cytokines (*e.g*. interleukin-6; IL-6) by limiting gut absorption and driving tissue sequestration of iron (*3*). Iron deficiency has been linked to impaired vaccine efficacy in humans and administration of a hepcidin mimetic limits CD8^+^ T cell responses to influenza vaccination and infection in murine models (*4, 5*). How systemic iron dysregulation shapes tumoral immunity has not been explored.

Pancreatic ductal adenocarcinoma (PDAC) has traditionally been considered immunologically ‘*cold*,’ characterized by an immunosuppressive tumor microenvironment (TME) with a relative paucity of infiltrating CD8^+^ T cells (TILs). Consistent with this, the response rate to immune checkpoint blockade (ICB) therapies in genomically unselected PDAC is < 5% (*6-8*). However, recent data have revealed significant immunological diversity in PDAC tumors, with increased CD8^+^ TILs correlating with improved disease outcomes in humans and ICB responsiveness in mice (*9, 10*). The factors underlying this immunological diversity remain unclear. As current evidence suggests PDAC has one of the highest rates of functional iron deficiency across solid tumors, we used PDAC as a model to explore the impact of systemic iron dysregulation on CD8^+^ T cell responses in cancer (*11, 12*). Through integrated analyses of human specimens and murine models of functional iron deficiency, we identify hepcidin-mediated iron sequestration as a novel metabolic checkpoint of anti-tumoral CD8^+^ T cell immunity.

### Functional iron deficiency is prevalent in PDAC and is associated with reduced survival

There are limited studies describing the prevalence and clinical impact of functional iron deficiency in patients with cancer. Therefore, we first performed a retrospective, pan-cancer analysis of institutional databases at Memorial Sloan Kettering Cancer Center (MSK) to investigate the prevalence of pre-treatment anemia (N=30,626) and iron deficiency (N=1,673) across patients with solid tumors. Given the increased prevalence of iron deficiency (serum transferrin saturation index [TSI] < 20%) in female patients (OR 1.6, 95% CI: 1.3 - 1.9, p < 0.0001), we focused on the ten most represented non-sex-restricted malignancies (N=1,332; Fig. 1A, Fig. S1A). While patients with luminal malignancies such as colorectal (CRC) and esophagogastric adenocarcinoma (EGC) had the highest rates of pre-treatment anemia (males: Hgb < 12 g/dL; females: Hgb < 11 g/dL) and iron deficiency, this was primarily absolute iron deficiency (serum ferritin ≤ 50 µg/L) likely due to chronic blood loss (Fig. 1A, Fig. S1A). Consistent with previous reports, patients with PDAC had one of the highest rates of functional iron deficiency (serum ferritin > 50 µg/L) across sex and disease stage (Fig. 1A,B; Fig. S1A-C) (*11, 12*). Unlike other malignancies, where functional iron deficiency was seen primarily in patients with advanced stage disease, functional iron deficiency was observed in PDAC patients across disease stages (Fig. 1B; Fig S1B). For patients with PDAC, pre-treatment anemia was associated with reduced overall survival (OS) independent of patient age, sex, and stage (Fig. 1C; Fig. S1D-F).

**Figure 1.**
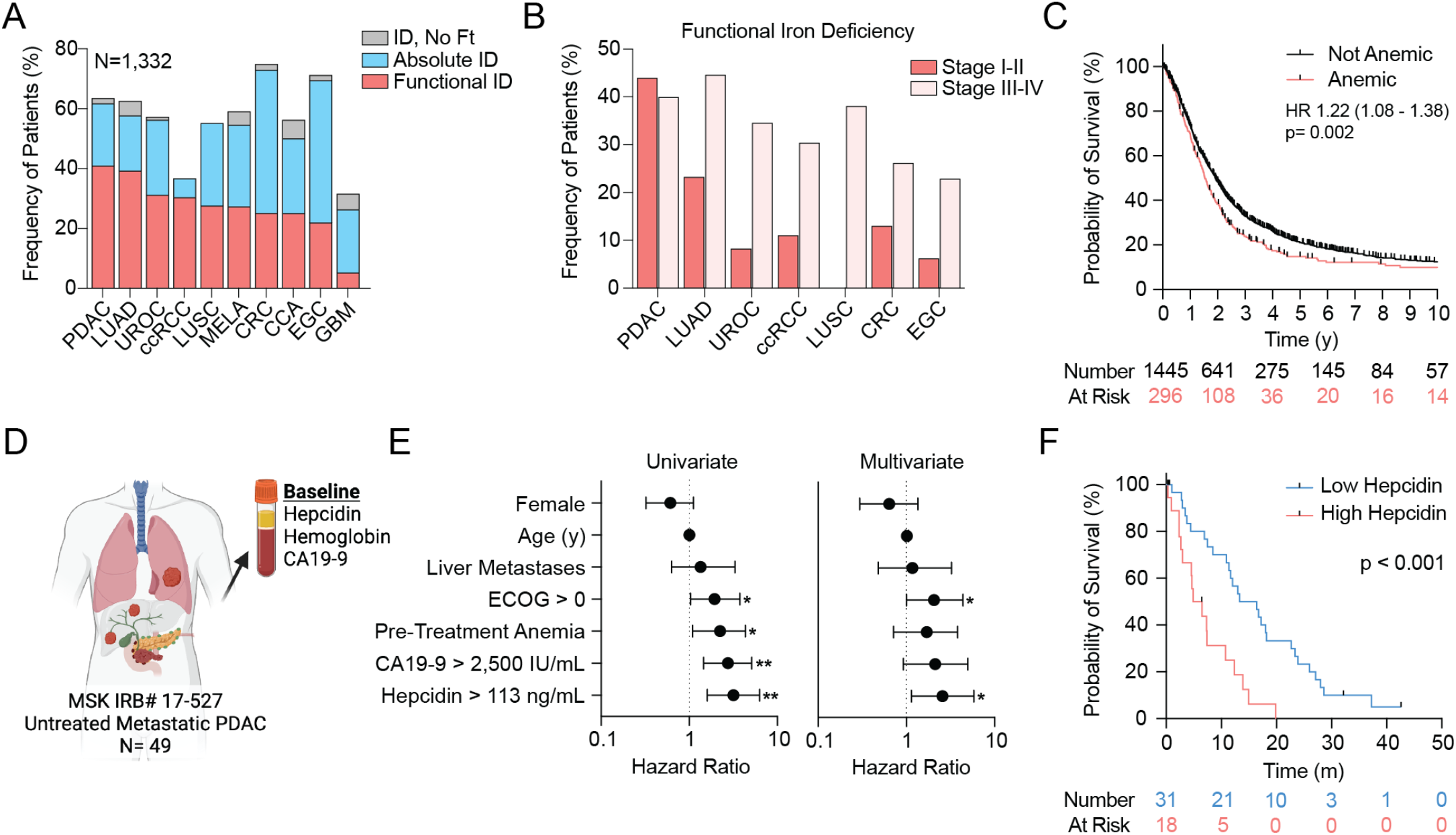
Functional iron deficiency is prevalent in PDAC and associated with reduced OS. (**A-B**) Prevalence of pre-treatment functional (TSI < 20%, ferritin > 50 µg/L) and absolute (TSI < 20%, ferritin ≤ 50 µg/L) iron deficiency (A) across patients with the 10 most-represented non-sex-restricted malignancies in MSK databases (N=1,332) and (B) stratified by disease stage (n ≥ 5 per stage group). PDAC: pancreatic ductal adenocarcinoma; LUAD: lung adenocarcinoma; UROC: urothelial carcinoma; ccRCC: clear cell renal cell carcinoma; LUSC: lung squamous cell carcinoma; MELA: melanoma; CRC: colorectal adenocarcinoma; CCA: cholangiocarcinoma; EGC: esophagogastric adenocarcinoma; GBM: glioblastoma multiforme. (**C**) Overall survival of patients with PDAC who presented with *vs*. without pre-treatment anemia analyzed by univariate Cox proportional hazard model. (**D-F**) Prospective cohort of patients with treatment naïve metastatic PDAC (N=49) enrolled on MSK #17-527. (D) Schema for sample collection. (E) Forest plot of univariate and multivariate Cox proportional hazard analysis of overall survival (hazard ratio [HR] ± 95% confidence intervals [CI]; *p<0.05; **p<0.01). (F) Kaplan-Meier survival curves of overall survival stratified by baseline serum hepcidin ≤ 113 ng/mL *vs*. > 113 ng/mL, p-value by log-rank test.

To specifically investigate the impact of functional iron deficiency on clinical outcomes, we next interrogated the association of baseline serum hepcidin levels with patient outcomes across a prospectively accrued cohort of patients (N=49) with newly diagnosed metastatic PDAC treated by physician choice (MSK #17-527; Fig. 1D; Fig. S1G). While performance status, pre-treatment anemia, elevated serum tumor marker carbohydrate antigen 19-9 (CA19-9), and elevated serum hepcidin were associated with reduced OS in univariate analyses, only performance status (HR 2.1, 95% CI: 1.0 – 4.4, p=0.049) and high serum hepcidin (HR 2.6, 95% CI: 1.4 - 5.8, p=0.023) remained independently associated with OS in multivariate analyses (Fig. 1E,F; Fig. S1H).

### Hepcidin limits the accumulation of CD8^+^ TILs

Hepcidin limits T cell responses to influenza vaccination and infection in murine models of functional iron deficiency (*5*). To investigate the relationship between functional iron deficiency and tumoral immunity, we analyzed pre-treatment peripheral blood and tumor samples from patients with newly diagnosed metastatic PDAC (N=28) treated with combination chemoimmunotherapy on the multi-institutional PRINCE trial (NCT03214250) (Fig. 2A) (*13*). Similar to the previous cohort, baseline serum hepcidin levels were independently associated with shorter OS (Fig. S2A,B). Integration of serum hepcidin levels with peripheral and intratumoral immune cell frequencies revealed a negative correlation between serum hepcidin and the frequency of intratumoral, but not circulating, CD8^+^ T cells (Fig. 2B; Fig. S2C) (*13*). Correlation of CD8^+^ TIL frequency with serum proteins revealed that hepcidin clustered alongside other known regulators of CD8^+^ T cell immunity and was independent of markers of either disease burden (CA19-9) or systemic inflammation (*e*.*g*. tissue necrosis factor (TNF), interferon-γ (IFN-γ)), indicating a unique role of hepcidin in regulating CD8^+^ T cell immunity in PDAC (Fig. 2C).

**Figure 2.**
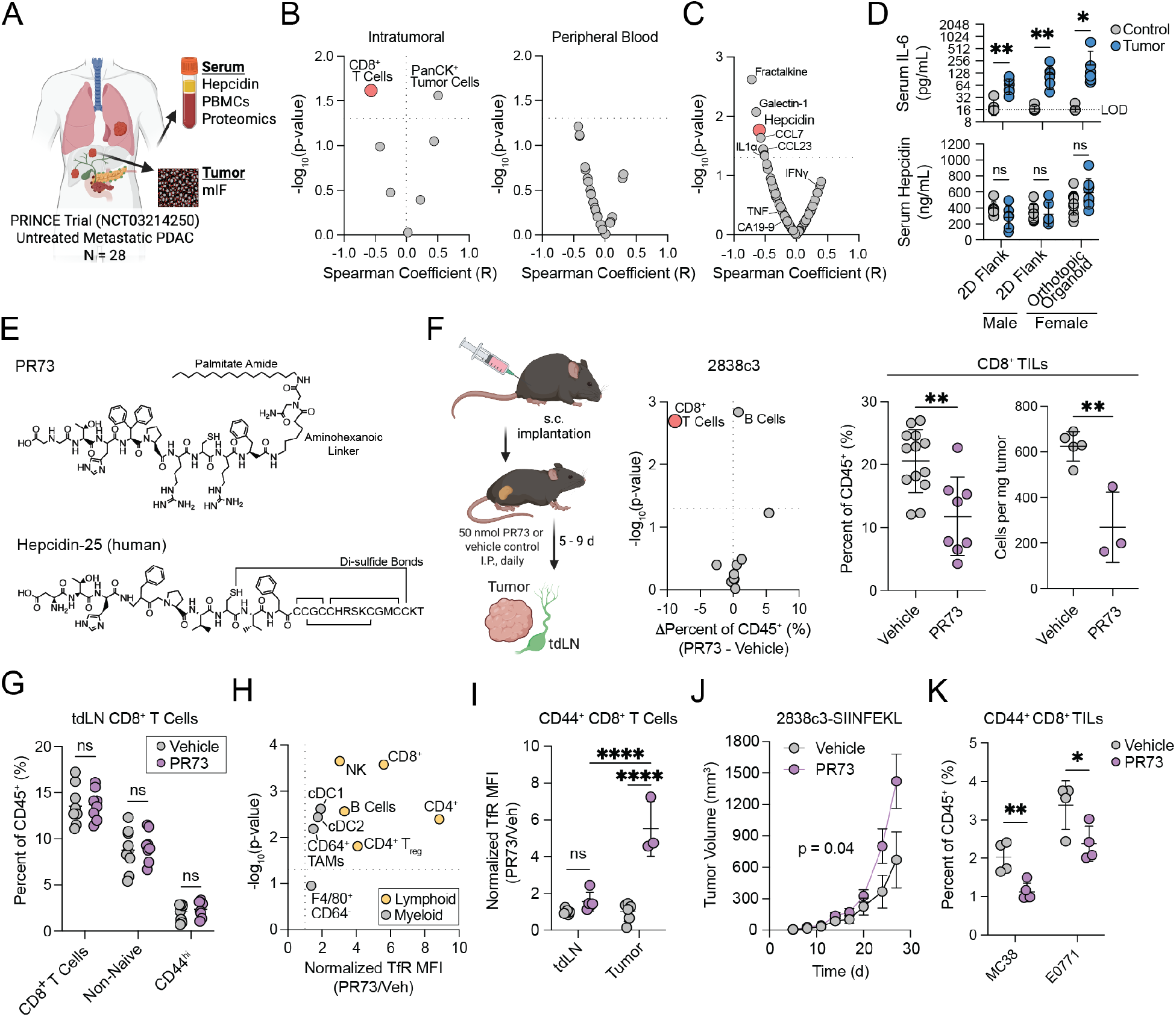
Hepcidin limits the accumulation of CD8^+^ TILs. (**A-C**) Patients with treatment-naïve metastatic PDAC enrolled on the PRINCE trial (NCT03214250). (A) Sample collection schema. (B) Spearman correlation between baseline serum hepcidin and the frequency of intratumoral or peripheral blood mononuclear cell subsets. (C) Spearman correlation between CD8^+^ TIL frequency and serum proteins. (**D**) Serum IL-6 and hepcidin from mice bearing 2838c3 flank tumors or orthotopic KPPC organoids *vs*. sex- and age-matched non-tumor-bearing controls. LOD = limit of detection (**E**) Chemical structure of PR73 and human hepcidin-25 (C-terminus: single letter amino acid code). (**F-I**) Schema and immunophenotype of 2838c3 (F) tumors and (G) tumor draining LN (tdLN) from PR73- or vehicle control-treated mice. Normalized TfR expression on (H) intratumoral immune cells and (I) CD44^+^ CD8^+^ T cells from tumors and tdLN. (**J**) Tumor growth (mean ± S.E.M.) of 2838c3-SIINFEKL tumors (25 nmol PR73, I.P. daily starting at day 5), p-value by mixed-effects model with Geisser-Greenhouse correction. (**K**) Frequency of CD44^+^ CD8^+^ TILs in vehicle and PR73-treated mice bearing MC38 (colon) and E0771 (breast) flank tumors. Unless otherwise noted, all error presented as mean ± S.D., pair-wise analyses by Student’s t-test, multiple comparisons by one-way ANOVA with Holm-Šidák correction (ns = not significant, *p<0.05, **p<0.01, ***p<0.001, ****p<0.0001).

To dissect the effect of hepcidin on CD8^+^ T cell immunity *in vivo*, we first asked whether conventional murine models of PDAC induce endogenous hepcidin expression. While implantation of KPPC (*LSL-Kras*^*G12D/+*^; *LSL-Trp53*^*R172H/R172H*^; *Pdx-1-Cre*)-derived PDAC organoids orthotopically in the pancreas or a KPC (*LSL-Trp53*^*R172H/+*^)-derived PDAC cell line (2838c3) in the flank reproducibly elevated serum concentrations of IL-6, serum hepcidin levels were unchanged when compared to age-matched non-tumor-bearing control mice (Fig. 2D) (*3, 14*). As nutritional iron contributes more to the circulating iron pool in mice than in humans, we hypothesized that elevated basal hepcidin expression driven by the high iron content of standard chow (184 ppm iron) might limit inflammation-driven hepcidin expression (*15*). However, hepcidin levels were unchanged in flank tumor bearing mice weaned on iron deficient (3 ppm), sufficient (39 ppm), or high iron (250 ppm) iso-nutritional diets compared to control non-tumor-bearing mice (Fig. S2E,F). Furthermore, while serum IL-6 correlated with hepcidin in PDAC patients (Fig. S2D), there was no correlation in tumor-bearing mice (Fig. S2G). These findings are consistent with reports in other tumor models that demonstrate that the tumor-driven hepatic hepcidin expression observed in patients is not recapitulated in many conventional mouse models of cancer (*16, 17*).

We therefore undertook a pharmacological approach to experimentally test the hypothesis that hepcidin-mediated iron sequestration causally restricts CD8^+^ TILs. Synthetic hepcidin mimetics with superior pharmacologic properties, such as PR73, have been developed as therapeutics for iron overload disorders (*18, 19*). The structure of PR73 resembles the first 9 amino acids of human hepcidin, which are essential for inhibition of the iron exporter ferroportin (Fpn), with the remaining 16 residues replaced by an aminohexonic linker and iminodiacetic palmitate amide, which has been suggested to enhance its binding affinity (Fig. 2E) (*18, 20*). As previously reported, administration of a single dose of PR73 (50 nmol, intraperitoneal (I.P.)) resulted in significant hypoferremia (Fig. S2H) (*5, 18*). Administration of PR73 (50 nmol/day, I.P.) for 9 days specifically limited the frequency and abundance of CD8^+^ TILs in a KPC-derived immunogenic model of PDAC (2838c3) without altering T cell subsets within the tumor-draining lymph node (tdLN) (Fig. 2F,G; Fig. S2I). Interestingly, PR73 most strongly induced transferrin receptor (TfR) expression on intratumoral lymphocytes as compared to intratumoral myeloid cells or CD44^+^ activated T cells within the tdLN (Fig. 2I, Fig. S2J).

KPC-derived murine models of PDAC are largely insensitive to anti-tumoral T cell immunity in the absence of enforced antigen expression (*21*). Expression of the MHC class I H-2K^b^-restricted antigen SIINFEKL in 2838c3 tumors (2838c3-SIINFEKL) significantly delayed tumor growth, consistent with antigen-specific anti-tumoral T cell immunity (Fig. S2K). Under these conditions, daily administration of PR73 (25 nmol, I.P.) significantly accelerated tumor growth and shortened survival as compared to vehicle-treated mice (Fig. 2J; Fig. S2L). Furthermore, PR73 also limited the frequency of activated CD8^+^ CD44^+^ TILs across implantable models of colon (MC38) and breast (E0771) carcinomas, supporting the generalizability of this phenomenon across malignancies (Fig. 2K).

### Hepcidin limits TIL iron availability

We next asked how PR73 limits intratumoral CD8^+^ T cell abundance. Addition of PR73 to culture medium did not alter the activation or growth of polyclonal splenic T cells *in vitro* (Fig. S3A). PR73 also did not alter T cell proliferation or expression of cell surface inhibitory receptors when cultured in the presence or absence of persistent TCR stimulation, suggesting an indirect mechanism by which PR73 limits CD8^+^ TILs *in vivo* (Fig. S3A). Furthermore, PR73 administration minimally impacted antigen-specific T cell priming in 2838c3-SIINFEKL-associated tdLNs, as characterized by intact upregulation of CD44 and downregulation of CD62L in adoptively transferred naïve antigen-specific OT-I^+^ CD8^+^ T cells (Fig. S3B).

We therefore hypothesized that PR73 limits CD8^+^ TILs primarily by altering iron availability within the TME. Consistent with this hypothesis, PR73 administration preferentially increased transferrin receptor (TfR) expression on TILs as compared to activated CD44^+^ CD8^+^ T lymphocytes in the tdLN (Fig. 2I). To quantify iron availability *in vivo*, we developed a ratiometric fluorescent reporter that exploits the endogenous iron responsive element/iron regulatory protein (IRE/IRP) system (Fig. 3A). The IRE/IRP system is a post-transcriptional regulatory network that maintains homeostatic levels of cytoplasmic labile iron. During iron scarcity, IRP1/2 bind IREs in the 5’ or 3’ untranslated region (UTR) of mRNA to block translation or increase mRNA stability of target transcripts, respectively (Fig. S3C). By placing the 5’ UTR of the mouse ferritin heavy chain (*Fth*) gene, containing a single IRE, upstream of mCherry, IRP binding blocks mCherry translation while the internal ribosome entry site (IRES) allows for ongoing GFP translation, resulting in an increase in the GFP/mCherry ratio (Fig. 3A).

**Figure 3.**
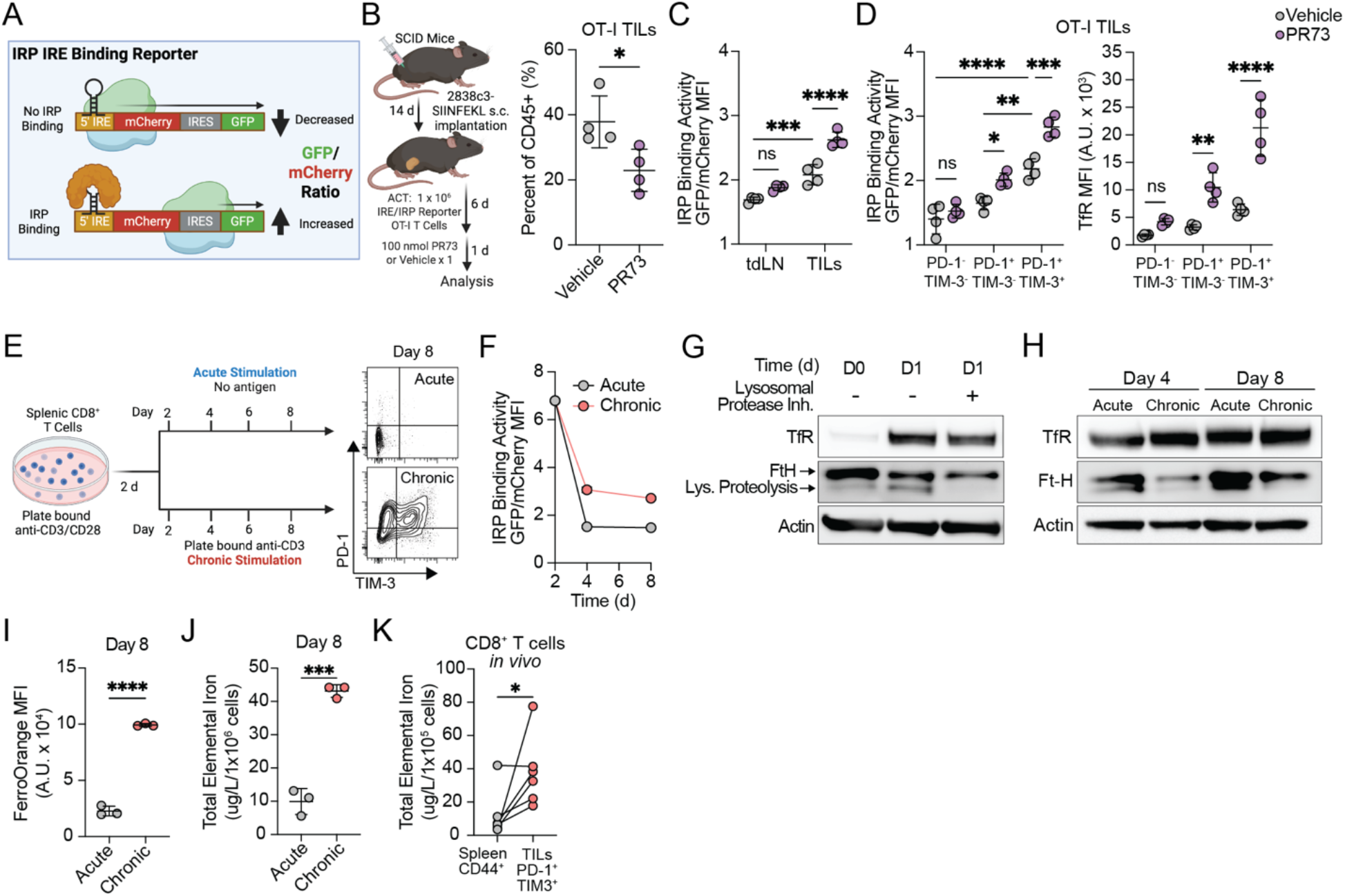
Persistent antigen stimulation disrupts CD8^+^ T cell iron metabolism. (**A**) IRP IRE binding reporter design. (**B-D**) (B) Experimental schema and frequency of OT-I TILs. (C-D) OT-I T cell IRP IRE binding activity or cell surface TfR expression in (C) the tdLN and tumor and (D) TILs, stratified by PD-1 and TIM-3 expression. (**E-I**) (E) *In vitro* platform of acute and chronic antigen stimulation. (F) IRP IRE binding activity. N-Ac (10 mM) added to culture medium from D2 to experimental endpoint. (G,H) Representative western blot of CD8^+^ T cells at specified timepoints. Lysosomal protease inhibitors E64d and pepstatin A (20 µg/mL) or DMSO control for 16h. (I) Labile Fe^2+^ as measured by FerroOrange. (**J**,**K**) Total elemental iron content of (J) *in vitro* cultured T cells or (K) FACS-sorted splenic CD44^+^ and intratumoral PD-1^+^ TIM-3^+^ CD8^+^ T cells as measured by graphite furnace atomic absorption spectrometry; p-value by paired Student’s t-test. All error presented as mean ± S.D., pair-wise analyses by Student’s t-test, multiple comparisons by one-way ANOVA with Holm-Šidák correction (ns = not significant, *p<0.05, **p<0.01, ***p<0.001, ****p<0.0001).

Following adoptive cell transfer (ACT) of activated IRE/IRP reporter expressing OT-I T cells into 2838c3-SIINFEKL tumor-bearing immunodeficient SCID hosts, mice were administered one dose of PR73 (100 nmol, I.P.) or vehicle control and paired tumors and tdLNs were isolated 24h later and assayed by flow cytometry (Fig. 3B). Antigen specific CD8^+^ TILs were reduced within 24 hours of PR73 administration suggesting an ongoing TIL requirement for iron within the TME (Fig. 3B). In vehicle-treated mice, IRE/IRP reporter activity was significantly higher in CD8^+^ TILs than CD8^+^ T cells within the tdLN suggesting basal levels of iron scarcity within the tumor (Fig. 3C). Furthermore, following PR73 administration, reporter activity was significantly increased in CD8^+^ TILs but not in CD8^+^ T cells within the tdLN (Fig. 3C). Both IRP activity and TfR expression were most significantly induced in terminally differentiated CD8^+^ TILs expressing both programmed cell death protein-1 and T-cell immunoglobulin and mucin-domain containing-3 (PD-1^+^ TIM-3^+^) (Fig. 3D).

### Persistent antigen stimulation disrupts CD8^+^ T cell iron metabolism

Tumor-infiltrating CD8^+^ T cells develop a hypofunctional cell state known as T cell ‘*exhaustion*’ that is driven primarily by chronic T cell receptor (TCR) signaling and marked by sustained expression of inhibitory cell surface receptors (*e*.*g*., PD-1, TIM-3) (*22, 23*). Given that PR73 induced IRP binding activity in TILs, but not in activated T cells within the tdLN (Fig. 3C), we hypothesized that hepcidin-mediated iron sequestration might impair T cell function specifically in the context of persistent antigen encounter. To isolate the impact of chronic TCR stimulation on iron demand, we first assessed cellular iron metabolism in activated polyclonal CD8^+^ T cells cultured in the presence or absence of persistent TCR stimulation *in vitro* (Fig. 3E). Consistent with previous reports, T cell iron demand increased following activation to support rapid cellular growth (*24, 25*), as evidenced by high IRP binding activity, rapid upregulation of iron uptake (TfR), and mobilization of stored iron from ferritin *via* lysosomal ferritinophagy (Fig. 3F,G). Following activation, and during cytokine-mediated expansion, acutely activated T cells re-established cellular iron homeostasis, characterized by reduced IRP binding activity, reduced TfR expression, and accumulation of ferritin (Fig. 3F,H). In contrast, persistent TCR stimulation disrupted cellular iron homeostasis resulting in sustained IRP binding activity and TfR expression despite increased cellular labile Fe^2+^, increased total elemental iron, and loss of the lower ferritin heavy chain band, consistent with reduced lysosomal ferritinophagy (Fig. 3F,H-J). A similar phenotype was observed *in vivo*, with isolated PD-1^+^ TIM-3^+^ CD8^+^ TILs containing significantly higher total elemental iron as compared to CD44^hi^ activated splenic CD8^+^ T cells (Fig. 3K).

The observation that persistent antigen stimulation sustained IRP binding activity despite elevated intracellular iron pools prompted us to hypothesize that chronic TCR stimulation disrupts functional iron utilization. The oxidation state of iron is critical to its function as an enzymatic co-factor; therefore, perturbations in the cellular redox state can disrupt the functional utilization of intracellular iron pools. For example, activation of IRP1 IRE binding by reactive oxygen species (ROS) requires disruption of the contained redox-sensitive iron-sulfur cluster (ISC), inducing a conformational change that promotes IRE binding (Fig. S3C) (*26*). Our group and others previously demonstrated that persistent antigen stimulation alone is sufficient to impair electron transport chain activity, resulting in increased steady-state levels of ROS and bioenergetic incapacitation of CD8^+^ T cells (*22, 23*). Therefore, we next tested whether persistent-antigen disrupts cellular iron metabolism *via* a redox-dependent mechanism.

Addition of the general thiol antioxidant and glutathione (GSH) precursor, N-acetylcysteine (N-AC, 10 mM), to chronically stimulated T cells is sufficient to restore mitochondrial electron transport and bioenergetic homeostasis (*22*). We found that N-AC completely abrogated persistent antigen-mediated increases in IRP activity and cytoplasmic labile Fe^2+^ (Fig. S3D,E). We therefore hypothesized that persistent antigen-driven mitochondrial dysfunction may broadly disrupt the function of other ISC-dependent processes, thereby sensitizing T cells to iron restriction only in the context of persistent antigen encounter. Consistent with this hypothesis, addition of a low concentration of the iron chelator desferrioxamine (DFO; 500 nM) to culture medium increased IRP binding activity, decreased the basal oxygen consumption rate (OCR) and mitochondrial membrane potential, and reduced proliferation of CD8^+^ T cells cultured in the presence of persistent antigen in a redox-dependent fashion (Fig. S3E-H).

### PR73 restricts the CD8^+^ TIL landscape

Intratumoral CD8^+^ T cell exhaustion is marked by progressive epigenetic and transcriptional rewiring as cells transition from a reversible to an irreversibly dysfunctional state (*27-33*). To identify T cell-intrinsic mechanisms regulating CD8^+^ TIL sensitivity to iron restriction, we analyzed adoptively transferred antigen-specific OT-I CD8^+^ TILs from vehicle or PR73-treated mice using integrated single-cell transcriptomic, epigenomic, and surface proteomic analysis (DOGMA-seq)(*34*). Unsupervised clustering and differential expression analysis of integrated transcriptomic and surface proteomic data led to the identification of four subsets including exhausted (Tex; *Cxcr6, Icos, Pdcd1, Havcr2*), proliferative exhausted (Tex-prolif; *Havcr2, Pdcd1, Mki67, Hells*), memory-like (Mem; *Cxcr3, Ccl5*), and effector-like (Eff; *Tnfrsf9, Ifng, Ccl4, Xcl1*) CD8^+^ TILs (Fig. 4A, Fig. S4A,B)(*35, 36*).

**Figure 4.**
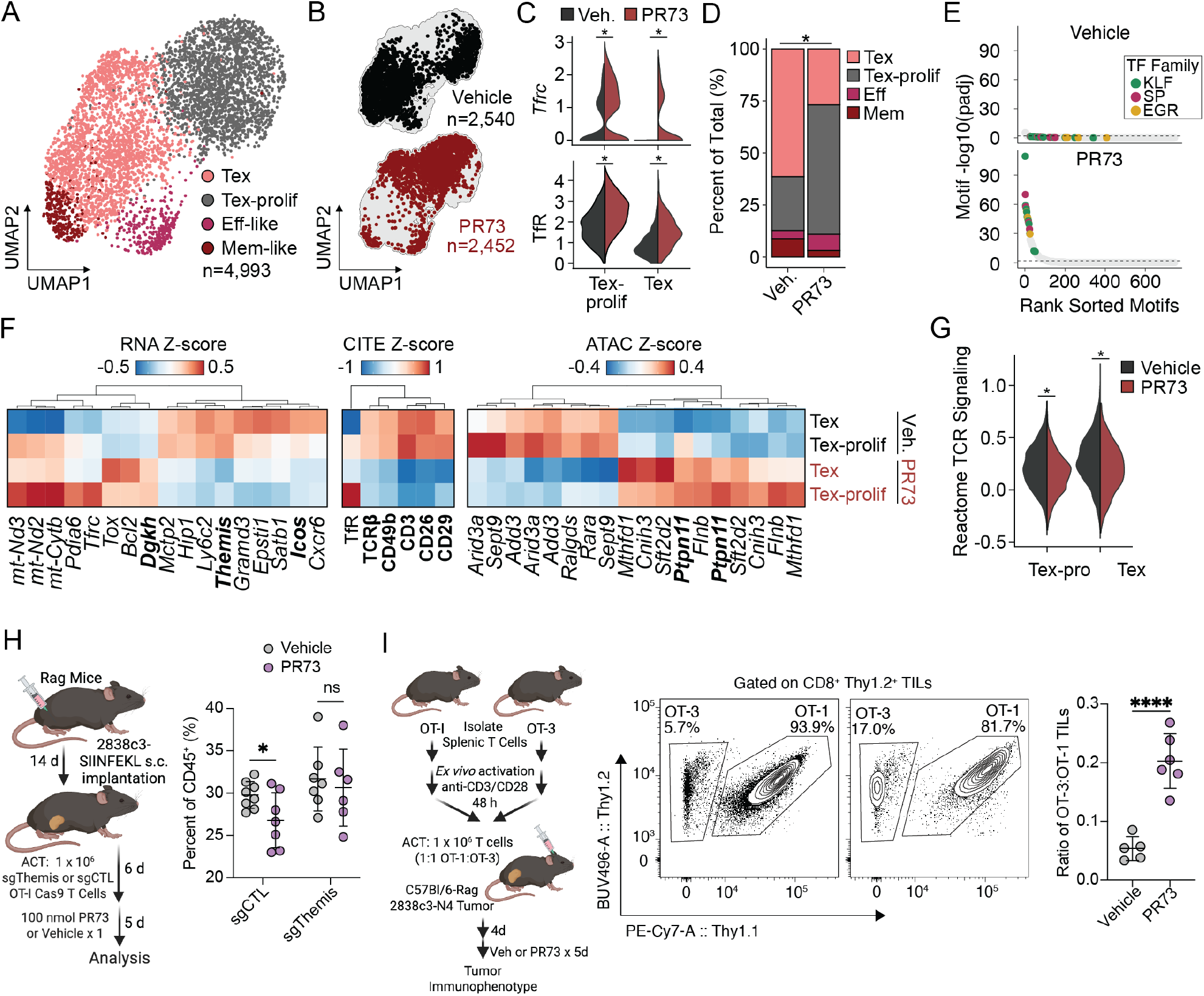
PR73 restricts the CD8^+^ TIL landscape. (**A-G**) Adoptively transferred OT-I TILs from vehicle and PR73 treated (50 nmol, I.P., daily for 7 days) 2838c3-SIINFEKL tumor-bearing mice were isolated and analyzed using the DOGMA-seq workflow. (A) Uniform manifold approximation and projection (UMAP) plot of integrated transcriptomic and surface proteomic data colored by cluster or (B) by treatment group. (C) *Tfrc* gene and TfR surface protein expression stratified by Tex subset and treatment. (D) TIL subtype proportions stratified by treatment (E) Differentially accessible transcription factor binding motifs plotted by rank-ordered adjusted p-values. (F) Z-score heatmaps for the top 5 differentially expressed genes, proteins, and peaks for each treatment group and Tex subtype. (G) REACTOME_TCR_SIGNALING module scores stratified by Tex subtype and treatment. (**H**) Frequency of *sgThemis* and *sgCTL* OT-I Cas9 TILs in vehicle and PR73-treated 2838c3-SIINFEKL tumor-bearing mice. (**I**) Representative flow cytometry plots and quantification of the ratio of OT-3:OT-1 TILs from vehicle and PR73-treated 2838c3-SIINFEKL tumor-bearing mice. All error presented as mean ± S.D. For flow cytometry, pair-wise analyses by Student’s t-test (ns = not significant, *p<0.05, **p<0.01, ***p<0.001, ****p<0.0001). Statistical analyses of DOGMAseq data are described in the Supplementary Methods.

All four subsets were identified across PR73 and vehicle-treated control mice (Fig. 4B). Consistent with our flow cytometry data (Fig. 3D), PR73 treatment significantly increased TfR expression at the RNA and protein levels in exhausted, but not memory- or effector-like, CD8^+^ TILs (Fig. 4C; Fig. S4C). PR73-administration resulted in a significant reduction in fractional Tex abundance, independent of cell cycle stage (Fig. 4D, Fig. S4D). Furthermore, accessible regions of chromatin in T cells isolated from PR73-treated mice were highly enriched for motifs bound by transcription factors known to be upregulated during iron deprivation, including early growth response (EGR) and krüppel-like factors/specificity protein (KLF/SP) families (Fig. 4E) (*37-39*).

Across modalities, we observed transcriptional (*Themis, Icos, Dgkh*), proteomic (CD3, TCRβ, CD49b/CD29, CD26), and chromatin accessibility (*Ptpn11*) changes associated with downregulation of TCR signaling (Fig. 4F) (*40-45*). More broadly, genes associated with TCR signaling (REACTOME_TCR_SIGNALING) were downregulated in exhausted CD8^+^ TILs from PR73-treated mice (Fig. 4G). Given our previous observations that persistent antigen stimulation is sufficient to disrupt mitochondrial function (*22*) and cellular iron metabolism (Fig. 3), we hypothesized that TCR signaling strength may calibrate the sensitivity of CD8^+^ T cells to iron restriction in the context of persistent TCR stimulation. To test this hypothesis, we leveraged several parallel genetic and pharmacological approaches to modulate TCR signal strength *in vitro* and *in vivo* (Fig. S4E). First, we cultured activated OT-I T cells in the presence of increasing concentrations of cognate antigen (SIINFEKL; N4) or in the presence of altered peptide ligands (APLs) with varying avidity for the OT-I TCR (N4>T4>V4)(*46*). Across models, persistent stimulation demonstrated antigen dose- and avidity-dependent increases in IRP binding activity, labile Fe^2+^, and TfR expression with associated decreases in lysosomal ferritinophagy (Fig. S4F,G). We confirmed the association of TCR avidity with iron demand by comparing OT-I transgenic T cells with OT-3 transgenic T cells, which express a rearranged TCR with approximately 100-fold lower avidity for N4 peptide (Fig. S4H) (*47*). Furthermore, OT-I T cells demonstrated reduced fitness during co-culture with OT-3 cells during persistent antigen stimulation in the presence of DFO (Fig. S4I).

Given that increased TCR signal strength sensitized CD8^+^ T cells to iron chelation *in vitro*, we hypothesized that attenuating TCR signaling would protect CD8^+^ TILs from intratumoral depletion during iron restriction *in vivo*. Among the most downregulated genes in exhausted TILs from PR73 treated mice was *Thymocyte-expressed molecule involved in selection (Themis*) (Fig. 4F), a known positive regulator of TCR signaling through the inhibition of Src homology phosphatase-1 (SHP-1), whose deletion results in increased phosphatase activity and reduced functional TCR avidity (Fig S4J)(*40, 41*). (*40, 41*). Indeed, CRISPR/Cas9-mediated deletion of *Themis* (*sgThemis*) rescued proliferation and reduced expression of both cell surface markers of T cell exhaustion and TfR during persistent antigen stimulation *in vitro* (Fig. S4K). We next adoptively transferred activated *sgThemis* or *sgCTL* OT-1 Cas9 T cells into 2838c3-SIINFEKL tumor-bearing immunodeficient mice. In contrast to *sgCTL* TILs, *sgThemis* TILs were resistant to PR73-mediated depletion (Fig. 4H).

These results suggested that PR73-mediated iron sequestration may remodel the repertoire of tumor-reactive TILs by selectively limiting CD8^+^ TILs responding to high avidity TCR-peptide major histocompatibility complex (pMHC) interactions *in vivo*. To directly test this hypothesis, we co-transferred *ex vivo* activated OT-I and OT-3 T cells into 2838c3-SIINFEKL tumor-bearing immunodeficient mice. Similar to *sgThemis* OT-I T cells *in vivo* as well as OT-3 cells co-cultured with OT-I T cells *in vitro*, intratumoral OT-3 T cells with reduced avidity for the N4 peptide were significantly protected from PR73-mediated deletion as compared to OT-I T cells *in vivo* (Fig. 4I). PR73 administration also induced significantly higher TfR expression in OT-I as compared to OT-3 TILs within the same tumor (Fig. S4L).

## Discussion

Systemic iron dysregulation has been an established hallmark of cancer for over fifty years, yet it remains unclear how systemic sequestration of an essential metabolite can promote tumor progression. Here, using PDAC as a model, we establish a fundamental role for hepcidin in limiting CD8^+^ T cell surveillance in cancer. Increased serum hepcidin is independently associated with reduced survival and inversely correlated with CD8^+^ TIL frequency in PDAC patients. Using a pharmacological model of murine functional iron deficiency to overcome species-specific differences in hepcidin regulation, we demonstrate that PR73-mediated depletion of CD8^+^ TILs occurs through a T-cell intrinsic, redox-dependent mechanism that is directly regulated by TCR signaling strength. This selective vulnerability preferentially restricts the accumulation of CD8^+^ TILs responding to high avidity interactions and suggests that systemic iron sequestration constrains the CD8^+^ TIL repertoire.

It is intriguing that a systemic metabolic perturbation was selectively disadvantageous to the fitness of intratumoral CD8^+^ T cells. Systemically, hepcidin drives iron sequestration primarily within macrophages of the reticuloendothelial system, a reflection of their role in erythrophagocytosis of senescent erythrocytes. Within the TME, both malignant and non-malignant cells with relatively higher iron avidity may further sequester interstitial iron. This has previously been shown to sustain mitochondrial function in tumor cells (*48*), but may further deplete iron availability to TILs. Whether targeting tumor- or myeloid cell-specific iron sequestration within the TME can improve the persistence of high-avidity CD8^+^ T cells within tumors is an exciting area of future exploration. The selective depletion of intratumoral, but not peripheral, CD8^+^ T cells also highlights the context-dependent nature of metabolic regulation and may explain why many tumor-reactive clones in patients are responding to low-avidity interactions. The preferential depletion of CD8^+^ T cells with high avidity for tumor-derived antigens may have broad implications for cancer immunotherapy and particularly for neoantigen vaccination strategies, which have shown early clinical promise in pancreas cancer but whose efficacy may depend on the induction of durable, high-avidity neoantigen-specific CD8^+^ T cell responses.

Identification of systemic iron sequestration as a key regulator of anti-tumoral immunity is notable given that it was identified in patients but not in traditional murine models of cancer. As a result, human translational studies will be required to determine whether targeting systemic iron sequestration is a viable therapeutic target in cancer. Identification of other environmental regulators of T cell immunity in patient samples, in combination with pharmacologic or genetic strategies to mimic not only the oncogenic drivers of tumorigenesis but also the unique disease-specific environments in mice, may further improve the biological and therapeutic relevance of these models.

## Acknowledgementss

We thank members of the Vardhana laboratory for discussion and critical feedback. We also acknowledge the MSK core facilities, including the Flow Cytometry Core Facility, Single Cell Analytics Innovation Lab and Integrated Genomics Operation, Cell Metabolism Core, and the Center of Comparative Medicine and Pathology. Finally, we thank the patients and their families for their participation.

## Author Contributions

J.D.S. and S.A.V. conceived the study. J.D.S. performed all experiments with assistance from J.J.N. and E.H. C.F. performed the pan-cancer analysis of pre-treatment anemia and functional iron deficiency with assistance and supervision from J.J., N.S., and Q.M. D.K. and K.H.Y. accrued and curated the cohort of patients with treatment naïve metastatic PDAC (MSK #17-527). J.E.C. curated the cohort of patients with treatment-naïve metastatic PDAC enrolled on the PRINCE trial. W.Z.Z. performed total elemental iron quantification with assistance and supervision from S.C. Single cell multiomics were performed and analyzed by C.S.M., R.C., and A.T.S. B.A. implanted orthotopic PDAC organoids with assistance and supervision from S.L. T.G. provided PR73 and T.G. and E.M.O. contributed additional work in conception, study guidance, and data analysis. J.D.S. and S.A.V. wrote the manuscript with assistance from C.S.M. All authors critically revised and approved the final manuscript.

## Data and materials availability

Raw multimodal scRNA-seq, ATAC-seq, and CITE-seq data will be deposited in GEO at the time of publication. Processed single-cell genomics data objects will be deposited into Synapse and companion analysis code for recreating main text Fig. 4 and supplemental Fig. S4 well be available on GitHub at the time of publication. Further information and requests for reagents may be directed to, and will be fulfilled by, the corresponding author(s).

## Supplementary Materials

Materials and Methods Figs. S1 to S6

