## Supplementary Materials for "Systemic iron sequestration restricts the intratumoral CD8^+^ T cell landscape in pancreas cancer"

### Materials and Methods

#### Human Studies

##### *Pan-cancer analyses of pre-treatment anemia and iron deficiency*

Patients treated at Memorial Sloan Kettering Cancer Center (MSK) were retrospectively identified through interrogation of institutional databases as approved by our Institutional Review Board (IRB). Specifically included were all patients with any of the top 15 most represented malignancies in MSK databases who previously consented to MSK IRB #12-245 (NCT01775072) and underwent paired tumor and normal next-generation sequencing *via* MSK-IMPACT (Integrated Mutation Profiling of Actionable Cancer Targets) prior to May 5, 2024 (N=32,982).

In descending order of representation and grouped by primary cancer type tumor registry annotations, these included: lung adenocarcinoma (LUAD), colorectal adenocarcinoma (CRC; colon adenocarcinoma, rectal adenocarcinoma, colorectal adenocarcinoma), invasive ductal carcinoma of the breast (IDC), prostate adenocarcinoma (PRAD), pancreatic adenocarcinoma (PDAC), urothelial carcinoma (UROC; bladder urothelial carcinoma, upper tract urothelial carcinoma), esophagogastric adenocarcinoma (EGC; stomach adenocarcinoma, esophageal adenocarcinoma, adenocarcinoma of the gastroesophageal junction, esophagogastric adenocarcinoma), high-grade serous ovarian cancer (HGSOC), uterine endometroid carcinoma (EEC), melanoma (MEL; cutaneous melanoma, melanoma), glioblastoma multiforme (GBM; glioblastoma multiforme, glioblastoma), cholangiocarcinoma (CCA; intrahepatic cholangiocarcinoma, cholangiocarcinoma, extrahepatic cholangiocarcinoma, perihilar cholangiocarcinoma), lung squamous cell carcinoma (LUSC), renal clear cell carcinoma (ccRCC), and invasive lobular carcinoma of the breast (ILC).

For each patient, the earliest three complete blood count (CBC), serum iron studies (iron, total iron binding capacity, transferrin saturation index), and ferritin lab values were extracted and then batch corrected *via* quantile matching, using the batch with the highest number of tests as the reference batch. Given the impact of anti-cancer therapies on blood counts and iron studies, only patients whose first lab tests were conducted prior to the prescription of any anti-cancer medication (chemotherapy, hormone therapy, targeted therapies, immunotherapies, biologics, anti-resorptive bone medications) within or outside MSK, as predicted by the MSK-CHORD (Clinicogenomic, Harmonized Oncologic Real-world Dataset) prior medication model applied to initial consult notes (*I*), were included (N=30,626 for CBC; N=1,673 for iron studies; N=1,935 for ferritin). For patients with > 1 pre-treatment lab value, only the earliest lab value was included in the final analyses. Anemia was defined as hemoglobin < 12 g/dL for male patients and < 11 g/dL for female patients. Iron deficiency was defined as a transferrin saturation index (TSI) < 20% and was further subtyped as absolute iron deficiency if serum ferritin was ≤ 50 ng/mL or functional iron deficiency if serum ferritin was > 50 ng/mL. For incidences of pre-treatment anemia and iron deficiency all patients with pre-treatment hemoglobin and TSI values, respectively, were included. Disease stage was defined as the patient's highest recorded stage (the maximum of AJCC clinical or pathologic staging) as of October 7, 2024.

We next evaluated the impact of pre-treatment anemia on survival in patients with PDAC (N=2,106). In addition to the presence or absence of pre-treatment anemia, patient's disease stage, gender, and age were extracted from the medical records. As patient's stage information is recorded at the time of primary cancer diagnosis and not at the time of anemia diagnosis, only patients whose stage diagnosis was < 60 days prior to pre-treatment labs were included (N=1,752). To limit immortality bias, data was left truncated, and patients removed whose death or date of last contact occurred prior to their earliest sequencing date (N=8). Using this cohort (N=1,744), left-truncated survival analyses comparing pre-treatment anemic vs. non-anemic patients were conducted using a univariate cox proportional hazard models (CoxPHFitter function in lifelines, with the sequencing date entered for the 'entry\_col' argument) across the cohort and within defined patient subsets.

#### Prospective cohorts of patients with untreated metastatic PDAC

Patients previously prospectively accrued on MSK IRB #17-527 (NCT03334708; "A study of blood-based biomarkers for pancreas adenocarcinoma") were retrospectively identified as having untreated metastatic pancreas adenocarcinoma (N=49). Clinical data was abstracted from the medical record. Anemia was defined as hemoglobin < 12 g/dL for male patients and < 11 g/dL for female patients. For CA19-9 non-secretors (CA19-9 < 1; N=1), CA19-9 was entered into the model as 0. Serum hepcidin was measured in baseline, pre-treatment serum samples by ELISA (R&D Systems, DHP250). The association of serum hepcidin and select clinical data were conducted by both univariate and multivariate cox proportional hazards regression. Overall survival was estimated using Kaplan-Meier methods and compared between patients with serum hepcidin above or below 113 ng/mL using log-rank test.

A second, independent cohort of patients (N=29) were previously prospectively accrued on the multi-institutional PRINCE trial (NCT03214250; "Safety and efficacy of APX005M with gemcitabine and nab-paclitaxel with or without nivolumab in patients with previously untreated metastatic pancreatic adenocarcinoma") (2). Quantification of baseline intratumoral cell frequencies by multiplex immunofluorescence and serum proteins by Olink® proteomics (Target 96 Immuno-oncology panel) were previously described and reported (2). For patients with > 1 quantified region of interest on multiplex immunofluorescence, the median across regions was used for correlative studies. Peripheral blood immune cells were assayed by CyTOF as previously described (2) and re-gated (Fig. S5). Serum hepcidin was measured in baseline, pre-treatment serum samples by ELISA (R&D Systems, DHP250). Overall survival was estimated using Kaplan-Meier methods and compared between patients with serum hepcidin above vs. equal to or below 21 ng/mL using log-rank test. Associations between immune cell subsets and serum markers were conducted by Spearman correlation.

#### **Animal Models**

##### Animals

All animal experiments were performed according to Memorial Sloan Kettering Cancer Center Institutional Animal Care and Use Committee guidelines (Protocol Number 20-10-012). Mice were purchased from The Jackson Laboratory: C57Bl/6J (#000664); C57Bl/6N (#005304); C57Bl/6J SCID (#001913); C57Bl/6J Rag1<sup>KO</sup> (#002216); C57Bl/6J OT-I (#003831); C57Bl/6 Rosa26-Cas9-EGFP (#026179). C57Bl/6J OT-3 were a gift from Dietmar Zehn (3). Heterozygous OT-1<sup>+/-</sup> Cas9-EGFP<sup>+/-</sup> mice were generated by breeding OT-1<sup>+/+</sup> with Cas9-EGFP<sup>+/-</sup> mice. Immunodeficient and TCR transgenic mice were implanted with tumors or sacrificed for T cell isolation, respectively, between 6-10 weeks of age. Immunocompetent mice were implanted with tumors or sacrificed for T cell isolation between 8-12 weeks of age.

#### *In vivo* tumor models

For flank tumor implantation in immunocompetent hosts, $2 \times 10^5$ tumor cells in 100 μL PBS were implanted subcutaneously in the right rear flank of 8-12-week-old syngeneic C57Bl/6J mice. For orthotopic implantation, PDAC organoids were dissociated into a single cell suspension and $2 \times 10^4$ single cells in 30 μL 50% PBS/Matrigel (Corning) were surgically implanted into the pancreatic tail of 12-week-old syngeneic C57Bl/6N mice as previously described (4). Unless specifically noted, all *in vivo* experiments were conducted in female mice weaned on standard chow (184 ppm iron; LabDiet PicoLab<sup>®</sup> Rodent Diet 20, #5053). For manipulation of nutritional iron, 4-week-old mice were provided iron deficient (3 ppm iron; TestDiet<sup>®</sup> 5WV4), iron sufficient (39 ppm iron; TestDiet<sup>®</sup> 58M1), or high iron (250 ppm iron; TestDiet<sup>®</sup> 5WV3) iso-nutritional diets (AIN-93M) until the experimental endpoint.

For analysis of adoptively transferred transgenic antigen-specific T cells in immunodeficient hosts, $2 \times 10^5$ 2838c3-SIINFEKL tumor cells in 100 μL PBS were implanted subcutaneously in the right rear flank of 8-10-week-old syngeneic C57BL/6J SCID or Rag1<sup>KO</sup> mice. Once tumors were established (~10 days), $1 \times 10^6$ naïve splenic OT-I<sup>+/-</sup> T cells, activated OT-I and OT-3 T cells (mixed 1:1), or transduced, activated splenic OT-I<sup>+/-</sup> Cas9-GFP<sup>+/-</sup> T cells (see “Primary mouse T cell isolation, activation, and culture” and “T cell retroviral transduction” below) in 200 μL PBS were adoptively transferred by retroorbital injection.

Tumor-bearing mice were monitored daily and euthanized when showing signs of morbidity as defined in the animal protocol. For experiments evaluating for tumor-associated induction of IL-6 and hepcidin, tumors were grown until the protocol-defined endpoint of tumor size or morbidity. For tumor immunophenotyping experiments, mice were sacrificed when median tumor volume reached approximately 500 μm<sup>3</sup>. Mice with tumor ulcerations were excluded as tumor ulceration was determined to alter systemic iron metabolism (data not shown). At the experimental endpoint, mice were euthanized with CO<sub>2</sub> and death was confirmed by cervical dislocation. Whole blood was collected by cardiac puncture, was allowed to clot for 30 – 60 min at room temperature (RT) and then spun at 2,700 x g for 10 min at RT. Serum was collected and frozen until analysis. Tissues were dissected, minced, and digested with 50 μg/mL Liberase TM (Roche), 100 μg/mL DNase I (Roche), and 350 units/mL hyaluronidase (Worthington) in RPMI media shaking at 250 rpm at 37°C for either 30 (spleen, tdLN) or 45 (tumor) minutes. Digested tissues were dissociated on a 100 μm cell strainer. For spleen, or if tissues were to be directly compared to spleen, red blood cells were lysed in ammonium-chloride-potassium (ACK) lysing buffer for 90 s and quenched with 10-fold volume of PBS. Single cell suspensions were then stained and analyzed by flow cytometry or subjected to FACS as outlined in “Flow Cytometry/FACS” below.

### Cell lines and organoids

Parental 2838c3 cell line, derived from a KPCY (*LSL-Kras*<sup>G12D/+</sup>; *LSL-Trp53*<sup>R172H/+</sup>; *Rose26-LSL-YFP*; *Pdx-1-Cre*) pancreatic tumor, was a gift from Dr. Katelyn Byrne (5). For all studies presented here, YFP was first deleted using lentiviral-mediated CRISPR-Cas9 (sgRNA: 5'-GTAGCCGAAGGTGGTCACGA-3') cloned into Lenti-CRISPRv2 plasmid (Addgene plasmid #52961) and selected by fluorescence-activated cell sorting (FACS). 2838c3-SIINFEKL cells were generated *via* retroviral transduction of PresentER-SIINFEKL-mCherry (Addgene plasmid #102945) into 2838c3 cells followed by purification by FACS. E0771 and MC38 cell lines were purchased from ATCC. Lenti-X<sup>TM</sup> cell line (Takara) was used for lentivirus and retrovirus generation and was maintained in DMEM + 10% FBS (GeminiBio; Benchmark<sup>TM</sup> #100-106) + 2 mM glutamine. All other cell lines were cultured in DMEM + 10% FBS. All cell lines were utilized prior to passage 20 and routinely checked by the MycoAlert Mycoplasma Detection Kit (Lonza; LT07-418) to be mycoplasma-negative. All cells were incubated at 21% O<sub>2</sub> in a humidity-controlled environment (37°C, 5% CO<sub>2</sub>; NuAire).

KPCY organoids were a kind gift of Dr. David Tuveson (4). Early passage organoids were cultured in complete organoid media with 10 μM Nutlin-3a (Sigma Aldrich) for three passages to select for cells that had undergone p53 loss-of-heterozygosity and then subsequently cultured under standard conditions as previously described (4).

### PR73 mini-hepcidin

For *in vivo* use, an appropriate mass of PR73 mini-hepcidin and Purebright SL220/Sunbright DSPE-020CN (NOF), at a ratio of 1 μmole:60 mg, respectively, were dissolved in 80% ethanol and dried in a Genevac EZ-2 Elite at RT until a gel is formed. The vehicle control solution was an equivalent mass of Purebright SL220 alone dissolved in 80% ethanol. The gel was re-dissolved in sterile de-ionized water to a final concentration of 50 nmoles of PR73 in 100 μL of water. Unless specifically indicated, mice were administered 100 μL of PR73 (50 nmol) or vehicle control daily by intraperitoneal (I.P.) injection. For *in vitro* experiments, PR73 was dissolved in DMSO and used at a final concentration of 50 nM.

### Primary Mouse T cell isolation, activation, and culture

Primary murine splenic T-cells were isolated by negative selection with the Dynabeads<sup>TM</sup> Untouched<sup>TM</sup> Mouse CD8<sup>+</sup> T Cells Kit (Invitrogen; 11417D) or Mouse T Cells Kit (Invitrogen; 11413D) following the manufacturer's instructions. Isolated T-cells were activated with plate-bound anti-CD3 antibody (clone 2C11, 3 μg/mL) and anti-CD28 antibody (clone 37.51, 1 μg/mL) in tissue culture treated 6-well plates for 2 days in complete T-cell media (RPMI-1640, 10% FBS, 2mM L-glutamine, 1X penicillin-streptomycin, 50 μM 2-mercaptoethanol, 10 ng/mL of murine IL-2 (Peprotech)) at a density of 1-1.5 x10<sup>6</sup> per mL as previously described (6).

For polyclonal CD8<sup>+</sup> T cells, T-cells are replated on day 2 in 6-well plates pre-coated with (chronic) or without (acute) anti-CD3 antibody (2C11, 3 μg/mL) at a density of 1x10<sup>6</sup> T-cell per ml in fresh complete T-cell media. For transgenic OT-I and OT-3 T cells, T-cells are replated on day 2 in 6-well plates in the presence (chronic) or absence (acute) of specified peptide (Biosynth) at a density of $1 \times 10^6$ T-cell per ml in fresh complete T-cell media. Cells are subsequently passaged similarly on days 4 and 6 at $1 \times 10^6$ and $1.5 \times 10^6$ (acute) and $1.5 \times 10^6$ and $2 \times 10^6$ (chronic), respectively. Where noted, DFO (500 nM, Sigma), N-Acetylcysteine (10 mM, Sigma), or PR73 (50 nM) were added to cell culture medium on days 2, 4, and 6. All cells were incubated at 21% O<sub>2</sub> in a humidity-controlled environment (37°C, 5% CO<sub>2</sub>; NuAire).

To determine the functional avidity of transgenic OT-I Cas9 CD8<sup>+</sup> T cells for SIINFEKL (N4) peptide, acutely activated T cells were restimulated on day 8 of culture with increasing concentrations of SIINFEKL peptide in fresh complete T-cell media at 37°C. After 1h, Brefeldin A (5 µg/mL) was spiked into the culture medium, and the cells were cultured for an additional 4 hours before cells were harvested and processed for flow cytometry.

#### IRP/IRE Reporter Cloning

The IRP/IRE binding reporter was designed using NEBuilder and cloned by Gibson Assembly into the mammalian retroviral expression vector pMIG (Addgene #9044) digested with EcoRI-HF (NEB; R3101) and PaeI (NEB; R0547). The 5' UTR of mouse *Fth* gene, containing a single IRE was PCR amplified from murine genomic DNA. Both mCherry-PEST (Addgene #176628) and EGFP-PEST (Addgene #172596) were used to shorten the fluorescent protein half-life, increasing dynamic reporter activity. The IRES was PCR amplified from the parental pMIG plasmid. Digested plasmid and PCR products were gel purified (Qiagen), assembled using Gibson Assembly 2X Mastermix according to the manufacturer's instructions (NEB E2611), and final plasmid sequence was confirmed *via* whole plasmid sequencing (Plasmidsaurus).

Gibson assembly PCR primers (Integrated DNA Technologies):

*mFTH\_5'UTR\_fwd*: 5'-TAGATCTCTCGAGGTTAACGGATCCCGCTATAAGTGCG-3'
*mFTH\_5'UTR\_rev*: 5'-TGCTCACCATCGGAACGACGAAGTTGCAAAG-3'
*mCherry-PEST\_fwd*: 5'-CGTCGTTCCGATGGTGAGCAAGGGCGAG-3'
*mCherry-PEST\_rev*: 5'-AGGGGGAATTTCACACATTGATCCTAGCAGAAG-3'
*IRES\_fwd*: 5'-CAATGTGTGAAATTCCCCCTAACGTTAC-3'
*IRES\_rev*: 5'-TGCTCACCATGGTTGTGGCCATATTATC-3'
*EGFP-PEST\_fwd*: 5'-GGCCACAACCATGGTGAGCAAGGGCGAG-3'
*EGFP-PEST\_rev*: 5'-AAGCGGTGATAATTCTTAATTTATACATTGATTCTCGCGC-3'

#### Lentivirus and retrovirus generation

Lentiviral and retroviral particles were generated using the Lenti-X<sup>TM</sup> cell line (Takara) with the Lipofectamine<sup>TM</sup> 3000 Transfection Reagent (Invitrogen) according to the manufacturer's instructions. The specified expression plasmid was co-transfected with packaging plasmids pMD2.G and psPAX2 (Addgene plasmids #12259 and #12260) for lentivirus generation or pCL-Eco (Addgene plasmid #12371) for retrovirus generation. Supernatants containing the viral particles were collected at 48 h and 72 h post transfection and were cleared of contaminating Lenti-X<sup>TM</sup> cells by centrifugation at 1200 rpm x 5 min. The supernatant was filtered through 0.45 µm polyethersulfone (PES) filters and concentrated with 8% polyethylene glycol (PEG) 8000

(Promega) overnight. Precipitated virus was concentrated 100x with fresh T cell media and frozen at -80C until use.

#### **Retroviral transduction of primary mouse T cells**

Transgenic splenic OT-I or OT-I Cas9 T cells were isolated by negative selection as described above (see “Primary Mouse T cell isolation, activation, and culture”) and activated on non-tissue culture-treated 6 well plates pre-coated with anti-CD3 and anti-CD28 overnight. The following day, non-tissue culture-treated 6-well plates were coated with 1.5 mL PBS with anti-CD3 antibody (clone 2C11, 3 µg/mL), anti-CD28 antibody (clone 37.51, 1 µg/mL), and RetroNectin® (Takara Bio, 13.3 µL/mL) for 90 min at 37C and then blocked with 2mL 2% bovine serum albumin (BSA) in PBS for 10 min at 37C. Following aspiration of the blocking solution, 2 mL of complete T cell media and 50 µL of retrovirus concentrate was added to each well and spun at 3000 rpm for 1 hour at 32C. Previously activated T cells were collected, spun at 350 x g for 8 min at 4C and resuspended in complete T cell media at a concentration of $1 \times 10^6$/mL. To the virus labeled plate, 2 mL of the T cell suspension was added on top of the virus media, and the plate was spun at 1200 rpm for 10 min at 4C and then cultured overnight at 37C. On day 2, transduction was confirmed by flow cytometry and cells were either cultured *in vitro* as described above or adoptively transferred into recipient immunodeficient mice.

#### **CRISPR-mediated gene editing of primary T cells**

Primary OT-I Cas9 splenic T cells were isolated by negative selection as described above (see “Primary Mouse T cell isolation, activation, and culture”) and retrovirally transduced with independent sgRNAs inserted *via* Gibson cloning into BbsI-HF (NEB) digested pMSCV-U6sgRNA(BbsI)-PGKpuro2ABFP (Addgene plasmid #102796) plasmid (see “Retroviral transduction of primary mouse T cells”). Knock-out of the gene of interest was confirmed by western blot.

*sgControl*: 5'-ACGCTATAGTGTACGTCTAA -3'

*sgThemis*: 5'-ACTGGATTTCTAGGACCCTG-3'

#### **Flow Cytometry/FACS**

When able, equivalent numbers of cells were stained across conditions. Single cell suspensions were centrifuged at 350 x g at 4C for 5 min and resuspended in Ghost Dye™ Viability Dye 510 or 780 (Tonbo) and Fc Block (Biolegend) for 10 minutes in PBS on ice. Cells were washed in FACS buffer (PBS + 2% FBS), centrifuged at 350 x g at 4C for 5 min and resuspended cell surface staining solution composed of cell surface marker antibodies at prespecified dilutions in 1:1 BD Horizon™ Brilliant Stain Buffer (BD Biosciences) and FACS buffer. Cells were stained for 30 (*in vitro* experiments) or 45 (*in vivo* experiments) minutes on ice, washed in FACS buffer, and centrifuged at 350 x g at 4C for 5 min.

For fluorescence-activated cell sorting (FACS), stained cells were resuspended in FACS buffer, filtered, and specified cell populations were sorted into FACS buffer using a 70 µm nozzle on either a FACS Aria™ III or FACSymphony™ S6 (BD Biosciences) flow cytometer in the MSK

Flow Cytometry Core Facility. For phenotyping experiments without intracellular staining, cell pellets were resuspended in 4% formaldehyde (Sigma) in PBS for 10 min at RT and then diluted in FACS buffer. For intracellular staining, cell pellets were fixed and permeabilized with the eBioscience™ FoxP3/Transcription Factor Fixation/Permeabilization Kit (Thermo Fisher) for 30 minutes at room temperature, washed with 1X Permeabilization Buffer, incubated with intracellular antibodies in 1X Permeabilization Buffer overnight at 4°C. The following day, permeabilized cells were washed with 1X Permeabilization Buffer, resuspended in 4% formaldehyde (Sigma) in PBS for 10 min at RT and then diluted in FACS buffer. Flow cytometry was performed on a Cytex® Aurora and data analysis was conducted using FlowJo™ v.10.10.0. Representative gating strategy for tumor immunophenotyping presented in Fig. S6.

Surface antibodies used are as follows: CD11b-BUV395 (BD Biosciences, clone M1/70, #563553, RRID: AB\_2738276), B220-BUV496 (BD Biosciences, RA3-6B2, 612950, AB\_2870227), Ly6G-BUV563 (BD Biosciences, 1A8, 612921, AB\_2870206), CD8a-BUV615 (BD Biosciences, 53-6.7, 613004, AB\_2870272), CD88-BUV661 (BD Biosciences, 20/70, 750080, AB\_2874295), CD44-BUV737 (BD Biosciences, IM7, 612799, AB\_287016), TCRb-BUV805 (BD Biosciences, H57-597, 748405, AB\_2872824), PD-L1-BV421 (Biolegend, 10F.9G2, 124315, AB\_10897097), CD45-BV480 (BD Biosciences, 30-F11, 566095, AB\_2872824), CD62L-BV570 (Biolegend, ME-14, 104433, AB\_10900262), F4/80-BV605 (Biolegend, BM8, 123133, AB\_2562305), Ly6C-BV650 (Biolegend, HK1.4, 128049, AB\_2800630), CD64-BV711 (Biolegend, X54-5/7.1, 139311, AB\_2563846), NK1.1-BV750 (BD Biosciences, PK136, 746876, AB\_2871676), XCR1-BV785 (Biolegend, ZET, 148225, AB\_2783119), CD39-PE Dazzle594 (Biolegend, Duha59, 143812, AB\_2750322), PD-1-PE/Cy7 (Biolegend, RMPI-30, 109110, AB\_572017), I-A/I-E-PE Fire810 (Biolegend, M5/114.15.2, 107667, AB\_289690), Ly108-APC (Biolegend, 330-AJ, 134609, AB\_2728154), CD4-AF700 (Biolegend, RM4-5, 100536, AB\_493701), CD11c-APC/Cy7 (Biolegends, N418, 117323, AB\_830646), CCR7-APC/Fire810 (Biolegends, 4B12, 120135, AB\_2910289), CD71-BUV395 (BD Biosciences, R17217, 567254, AB\_2916516), CD62L-BUV737 (BD Biosciences, MEL-14, 612833, AB\_2870155), TIM3-BV421 (Biolegends, RMT3-23, 119723, AB\_2616908), CD44-BB700, (BD Biosciences, IM7, 566506, AB\_2870126), PD-1-APC/Cy7 (Biolegends, 29F.1A12, 135224, AB\_2563523), CD45-BV570 (Biolegend, 30-F11, 103136, AB\_2562612), CD11b-BV785 (Biolegend, M1/70, 101243, AB\_2561373), TIM3-PE (Biolegend, RMT3-23, 119704, AB\_345378), PD-1-APC (Biolegend, RPMI-30, 109112, AB\_10612938), Thy1.2-BUV805 (BD Biosciences, 53-2.1, 741908, AB\_2871222), Thy1.1 (Biolegend, OX-7, 202518, AB\_1659223). The intracellular antibodies used are as follows: TCF1/7-AF488 (BD Biosciences, S33-966, 560718, AB\_2916388), TCF1/7-A647 (BD Biosciences, S33-966, 566693, AB\_2869823), FoxP3-PE Cy5 (Invitrogen, FJK-16s, 15-5773-82, AB\_468806), IFNγ-PE-Cy7 (Biolegend, XMG1.2, 505826, AB\_2295770), TNF-BV650 (Biolegend, MP6-X).

### Immunoblotting

Single cell suspension cells were centrifuged at 500 x g at 4°C for 5min, washed in cold PBS, and pelleted at 700 x g for 2 min at 4°C. Cell pellets were lysed in cold RIPA buffer (Cell Signaling, #9806) supplemented with HALT™ protease and phosphatase inhibitor cocktails (Thermo Fisher) for 30 min on ice. Lysates underwent three freeze-thaw cycles, and cellular debris was removed by centrifugation at 20,000 x g for 20 min at 4°C. Protein concentration of collected supernatants were quantified using the Pierce<sup>TM</sup> BCA Protein Assay Kit (Thermo Fisher) following the microplate protocol on a SpectroStar nano. An appropriate volume of sample to load 10-25 µg of protein per well was mixed with NuPage<sup>TM</sup> 4X LDS Sample Buffer and NuPage<sup>TM</sup> 10X Sample Reducing Buffer (Invitrogen), heated for 10 min at 70°C, and separated on NuPage<sup>TM</sup> Bis-Tris 4-12% gels at 100V for 135 min in NuPage<sup>TM</sup> MOPS SDS running buffer. Proteins were transferred to a 0.45 µm nitrocellulose membrane at 100 V for 75 min in NuPage<sup>TM</sup> transfer buffer containing 20% methanol. Membranes were blocked with 5% skim milk (BD Difco<sup>TM</sup>) in TBST (TBS + 0.1% Tween) for 1hr at RT, washed in TBST (3 x 5 min), and incubated in primary antibody overnight at 4°C in TBST + 3% BSA. Primary antibodies: anti-Transferrin Receptor/TfR (1:1,000, RRID: AB\_2533029), anti-Ferritin Heavy Chain/FtH (1:1000, RRID: AB\_1310222), and anti-Actin (1:5,000, RRID: AB\_476697). Blots were washed with TBST (3 x 5 min) and incubated with HRP-conjugated secondary antibodies (1:5000 in 5% milk in TBST) for 1hr at RT. Secondary antibodies: horse anti-mouse (RRID: AB\_330924), goat anti-rabbit IgG (RRID: AB\_228341). Membranes were washed with TBST (3 x 5 min), incubated with 1-Shot Digital-ECL Solution (Kindle Biosciences) or Pierce<sup>TM</sup> ECL Western Blotting Substrate (Thermo Scientific) for 1 min at RT, and imaged on a Bio-Rad ChemiDoc<sup>TM</sup> Touch Gel Imaging System.

#### **Total elemental iron quantification**

Isolated T cell samples were washed twice in PBS, dry pelleted in low-iron tubes, and frozen. Samples are processed by adding a digestion buffer, 50% nitric acid (v/v), to the sample in a 1:1 ratio and then heated at 70°C for 2 hours in a heat block. The digested samples were then removed, cooled to room temperature, and centrifuged for 5 minutes at 6000 x g. The supernatant was collected and then diluted in dilute 0.2% nitric acid for measurement via the graphite furnace atomic absorption spectrometer (PerkinElmer model 900z). Iron was measured in the digestion buffer to assess iron contamination, and sample iron levels were quantified using a standard curve of known iron concentrations and normalized to cell number.

#### **Single cell multiomics (DOGMA-seq)**

##### *Sample Preparation, Library Construction, and Next-Generation Sequencing*

Following establishment of 2838c3-SIINFEKL tumors in C57Bl/6J Rag1<sup>KO</sup> mice, mice were administered PR73 (50 nmol, I.P., daily) or vehicle control for 2 days prior to the adoptive transfer of 1 x 10<sup>6</sup> naïve antigen-specific OT-I T cells. Following treatment with PR73 or vehicle control for an additional 7 days, tumors were dissociated, and OT-I TILs were isolated by FACS as described above. In addition to fluorescently tagged antibodies for FACS, single cell suspensions were labeled with TotalSeq-A Hash Tag Oligonucleotide (HTO)-conjugated antibodies (7) to allow for sample multiplexing (1:100, Biolegend) and TotalSeq-A Universal Cocktail of Antibody-Derived Tags (ADTs) to enable simultaneous detection of surface protein expression of selected extracellular epitopes, following the manufacturer's protocol (Biolegend). Following FACS, HTO-tagged samples were combined and processed as a single, multiplexed sample.

Single-cell Multiome ATAC + Gene Expression was performed using the Chromium Next GEM Single Cell Multiome system (10x Genomics) according to the manufacturer's recommendations, with modifications to the single-cell preparation step. Specifically, sample preparation was adapted from the DOGMA-seq protocol described by Mimitou et al. (8). Briefly, cells were washed once in PBS (without $\text{Ca}^{2+}/\text{Mg}^{2+}$) and resuspended at $\sim 1 \times 10^6$ cells/ml. The suspension was incubated with digitonin lysis buffer composed of 20 mM Tris-HCl (pH 7.4), 150 mM NaCl, 3 mM $\text{MgCl}_2$, 0.01% digitonin (DIG; MilliporeSigma), and 2 U $\mu\text{l}^{-1}$ RNase inhibitor (Promega). Lysis was performed on ice for 5 min. Following lysis, 1 ml of ice-cold DIG wash buffer (20 mM Tris-HCl, pH 7.4; 150 mM NaCl; 3 mM $\text{MgCl}_2$; and 1 U $\mu\text{l}^{-1}$ RNase inhibitor) was added to quench the reaction. The suspension was gently inverted several times and centrifuged at $500 \times g$ for 5 min at 4 °C. The supernatant was carefully removed, and the cell pellet was resuspended in 100 $\mu\text{l}$ of chilled DIG wash buffer. Cells were counted using trypan blue on a Countess II FL Automated Cell Counter (Thermo Fisher Scientific) and diluted to the desired concentration to capture a target recovery of 10,000 cells per sample.

The cell suspension was loaded into a Chromium Next GEM Chip J, and partitioning into Gel Bead-in-Emulsions (GEMs) was performed using the Chromium Controller (10x Genomics). Library construction for both ATAC and gene expression (GEX) libraries was carried out following the Chromium Next GEM Single Cell Multiome ATAC + Gene Expression Reagent Kits User Guide (10x Genomics, CG000338), including transposition, reverse transcription, cDNA amplification, and library indexing. ADTs and HTOs were co-processed with the Multiome libraries, enabling integrated quantification of ATAC, GEX, HTO and ADT modalities from the same cells. Final ATAC, GEX, HTO, and ADT libraries were pooled according to 10x Genomics recommendations (based on expected read depth per modality) and sequenced on an Illumina NovaSeq X Plus platform using paired-end sequencing. scRNA-seq, scATAC-seq, and CITE-seq libraries were sequenced to an average of 26,074, 34,627, and 7,784 reads per cell, respectively.

##### Data Pre-Processing and Quality-Control

scRNA-seq, scATAC-seq, and CITE-seq library FASTQs were pre-processed using Cell Ranger ARC Version 2.0.2 (10x Genomics) and aligned to the GRCm38 (mm10) reference genome. Antibody-derived tag (ADT) reads were aligned to the TotalSeq-A Universal Cocktail reference (Biolegend). Filtered Cell Ranger outputs were used to generate a multimodal analysis object using the Seurat (9) and Signac (10) R packages. Sample demultiplexing based on hashtag oligonucleotide counts was performed using the deMULTIplex2 R package (11) before multiplets, unclassified cells, and low-quality clusters (i.e., doublet-enriched clusters and/or clusters with low RNA or ATAC counts) were removed as described previously (12). Cleaned data was then re-processed using Seurat and Signac and then used for unsupervised clustering, differential gene expression testing, manual annotation of major TIL subtypes, motif analysis, and differential TIL subtype proportion testing as described in the Results section.

##### Statistical Test and Visualizations

TIL subtype annotation makers were identified using the 'FindAllMarkers' function applied to the scRNA-seq data, and the top 8 statistically significant marker genes were plotted using the

‘DotPlot’ function in Seurat. Statistically significant differences between PR73- and vehicle-treated exhausted TILs were identified using the Wilcoxon rank-sum test as implemented in the ‘FindMarkers’ function and visualized using the ‘VlnPlot’ function in Seurat. Differentially accessible transcription factor binding motifs were identified using the ‘FindMotifs’ function applied to the top 100 differentially accessible peaks between PR73- and vehicle-treated exhausted TILs identified using the ‘FindMarkers’ function applied to the scATAC-seq data. Differentially accessible motifs were plotted by rank-ordered Bonferroni-adjusted p-values. Statistically significant shifts in TIL subtype proportions were identified using the ‘propeller’ function with bootstrapping in the ‘Speckle’ R package (13). Treatment-specific marker gene z-score heatmaps were computed using the top 5 differentially expressed genes, proteins, and peaks for each treatment group and exhausted TIL subtype (identified using the ‘FindMarkers’ function) and visualized using ComplexHeatmap. Peaks included in the scATAC-seq z-score heatmap were annotated according to the closest feature as determined using the ‘ClosestFeature’ function in Signac. REACTOME\_TCR\_SIGNALING module scores were computed by first subsetting the REACTOME\_TCR\_SIGNALING gene list to include only those in the top 2,000 variably expressed genes in the dataset before module computation using the ‘AddModule’ function in Seurat. All visualizations were generated using the Seurat, ggplot2 (14), and ComplexHeatmap (15) R packages.

### Statistics

Unless otherwise noted above (‘Pan-cancer analyses of pre-treatment anemia and iron deficiency’ and ‘Single cell multiomics (DOGMA-seq)’), GraphPad PRISM 10 software was used for all statistical analyses. Error bars, p-values, and statistical tests are reported in figure legends.

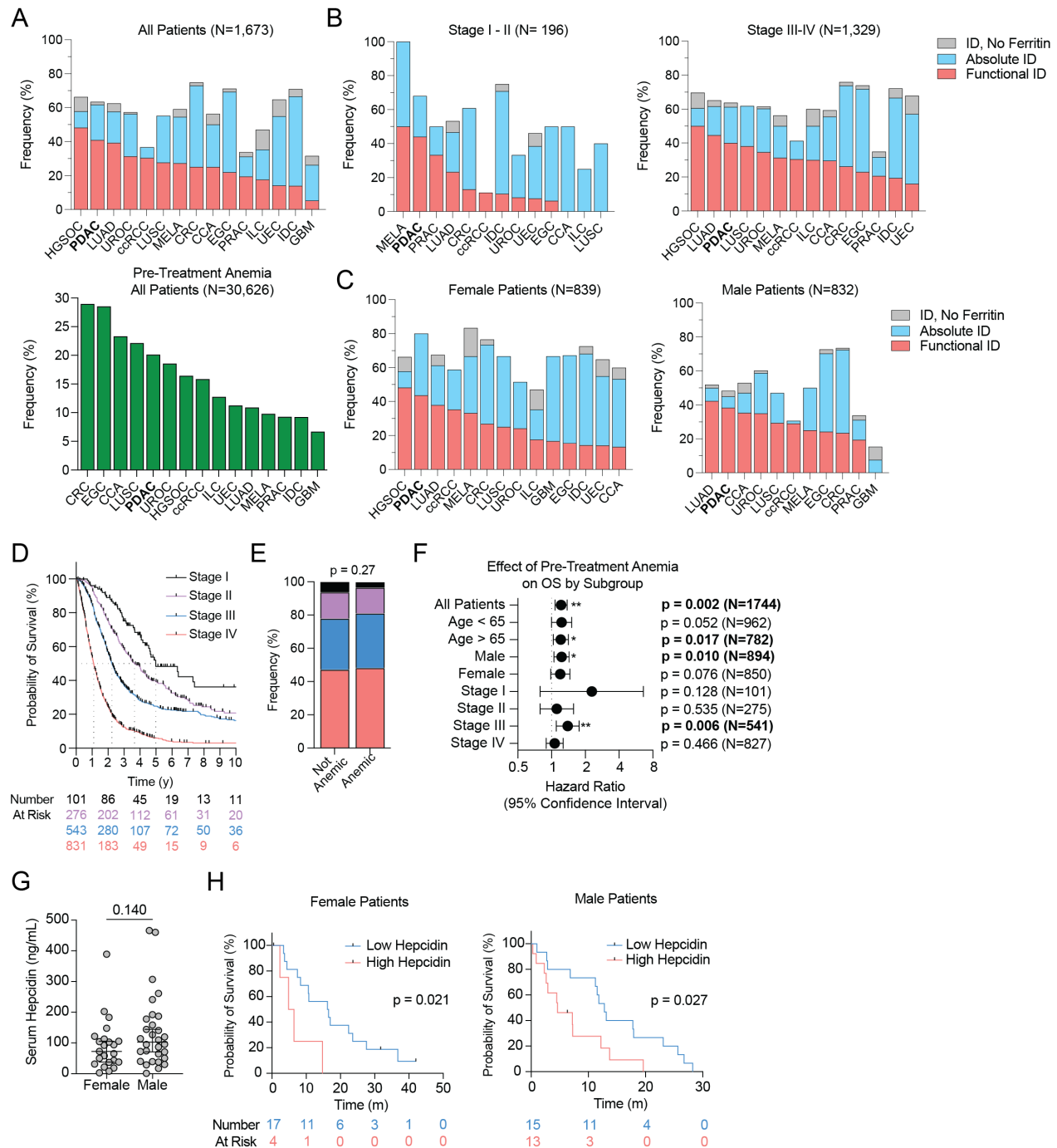

**Fig. S1. Related to Figure 1. Pan-cancer prevalence of pre-treatment anemia and iron deficiency.** (A-C) Prevalence of pre-treatment anemia (males: Hgb < 12 g/dL; females: Hgb < 11 g/dL) and functional (TSI < 20%, ferritin > 50 μg/L) and absolute (TSI < 20%, ferritin ≤ 50 μg/L) iron deficiency (ID) (A) across patients with the 15 most-represented malignancies in MSK databases and (B) stratified by disease stage and (C) gender. HGSOc: high-grade serous ovarian cancer; PDAC: pancreatic ductal adenocarcinoma; LUAD: lung adenocarcinoma; UROC: urothelial carcinoma; ccRCC: clear cell renal cell carcinoma; LUSC: lung squamous cell carcinoma; MELA: melanoma; CRC: colorectal adenocarcinoma; CCA: cholangiocarcinoma;

EGC: esophagogastric adenocarcinoma; PRAC: prostate adenocarcinoma; ILC: invasive lobular carcinoma of the breast; UEC: uterine endometroid carcinoma; IDC: invasive ductal carcinoma of the breast; GBM: glioblastoma multiforme. **(D-F)** Patients with PDAC and pre-treatment hemoglobin concentrations across MSK databases. **(D)** Kaplan-Meier curves of OS stratified by disease stage. **(E)** Stage distribution by pre-treatment anemia; p-value by Fisher's exact test. **(F)** Overall survival of patients with *vs.* without pre-treatment anemia analyzed by Cox proportional hazards models stratified by patient age, sex, and stage. **(G,H)** Prospective cohort of patients with treatment-naïve metastatic PDAC (N=49) enrolled on MSK #17-527. **(G)** Baseline serum hepcidin concentrations across patient sex with median ± 95% CI; p-value by Mann-Whitney test. **(H)** Kaplan-Meier survival curves of overall survival stratified by baseline serum hepcidin ≤ 113 ng/mL *vs.* $>$ 113 ng/mL; p-value by log-rank test.

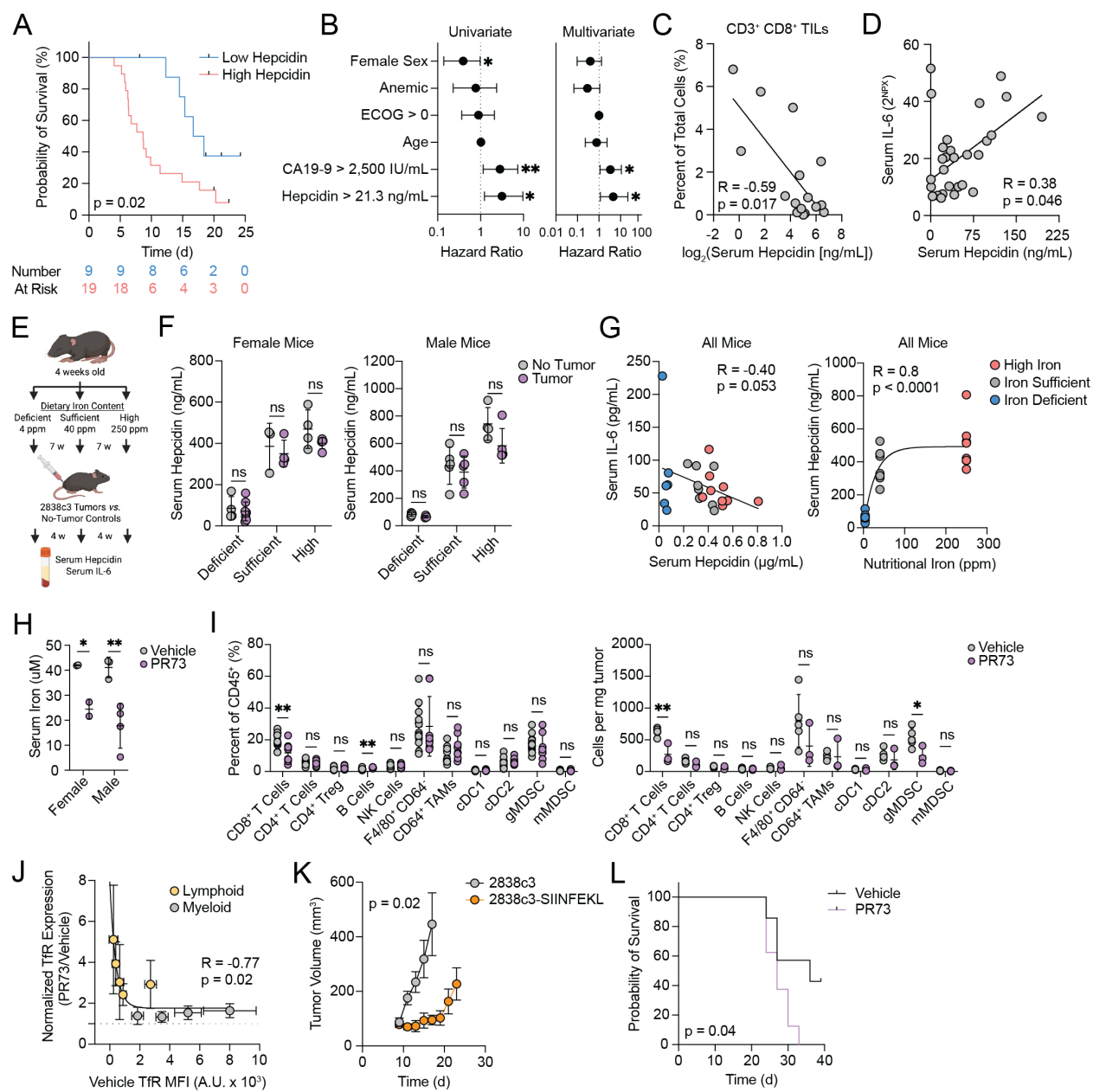

**Fig. S2. Related to Figure 2. Hepcidin limits CD8<sup>+</sup> T cell accumulation and function in patients and murine models of PDAC.** (A-D) Prospective cohort of patients with treatment-naïve metastatic PDAC (N=28) enrolled on the PRINCE trial (NCT03214250). (A) Kaplan-Meier curves of OS stratified by baseline serum hepcidin $\leq 21.3$ ng/mL vs. $> 21.3$ ng/mL; $p$-value by log-rank test. (B) Forest plot of univariate and multivariate Cox proportional hazard analyses of OS (HR ± 95% CI). (C-D) Spearman correlation of baseline serum hepcidin and (C) CD8<sup>+</sup> TILs and (D) serum IL-6 (O-link proteomics, NPX = normalized protein expression); least-squares regression fit shown for visualization. Each dot represents an individual patient. (E-G) C57Bl/6 mice weaned onto iso-nutritional chow containing varying dietary iron at 4 weeks of age. (E) Experimental schema. (F) Serum hepcidin in tumor-bearing and sex- and age-matched non-tumor-bearing control mice. (G) Spearman correlations between serum hepcidin and serum

IL-6 or dietary iron content; least-squares regression fit for visualization. **(H)** Serum iron from 8-week-old C57Bl/6 mice 16h after intraperitoneal (I.P.) administration of 50 nmol PR73 or vehicle control. **(I)** Frequency and abundance of intratumoral immune cell subsets following administration of 50 nmol PR73 or vehicle control I.P. daily for 5-9 days. **(J)** Spearman correlation of TfR expression per intratumoral cell subset in vehicle-treated mice and normalized PR73-mediated induction of TfR. **(K)** Volume (mean ± S.E.M.) of 2838c3 and 2838c3-SIINFEKL flank tumors, p-value by mixed-effects model with Geisser-Greenhouse correction. **(L)** Kaplan-Meier survival curves of mice bearing 2838c3-SIINFEKL tumors treated with PR73 (25 nmol, I.P. daily starting on day 5); endpoint defined as tumor volume $\geq 1000 \text{ mm}^3$; p-value by log-rank test. Across experiments, each point represents an individual patient or mouse except for panels J and K. Unless otherwise noted, data presented as mean ± S.D., pair-wise analyses by Student's t-test. \*p<0.05; \*\*p<0.01; \*\*\*p<0.001.

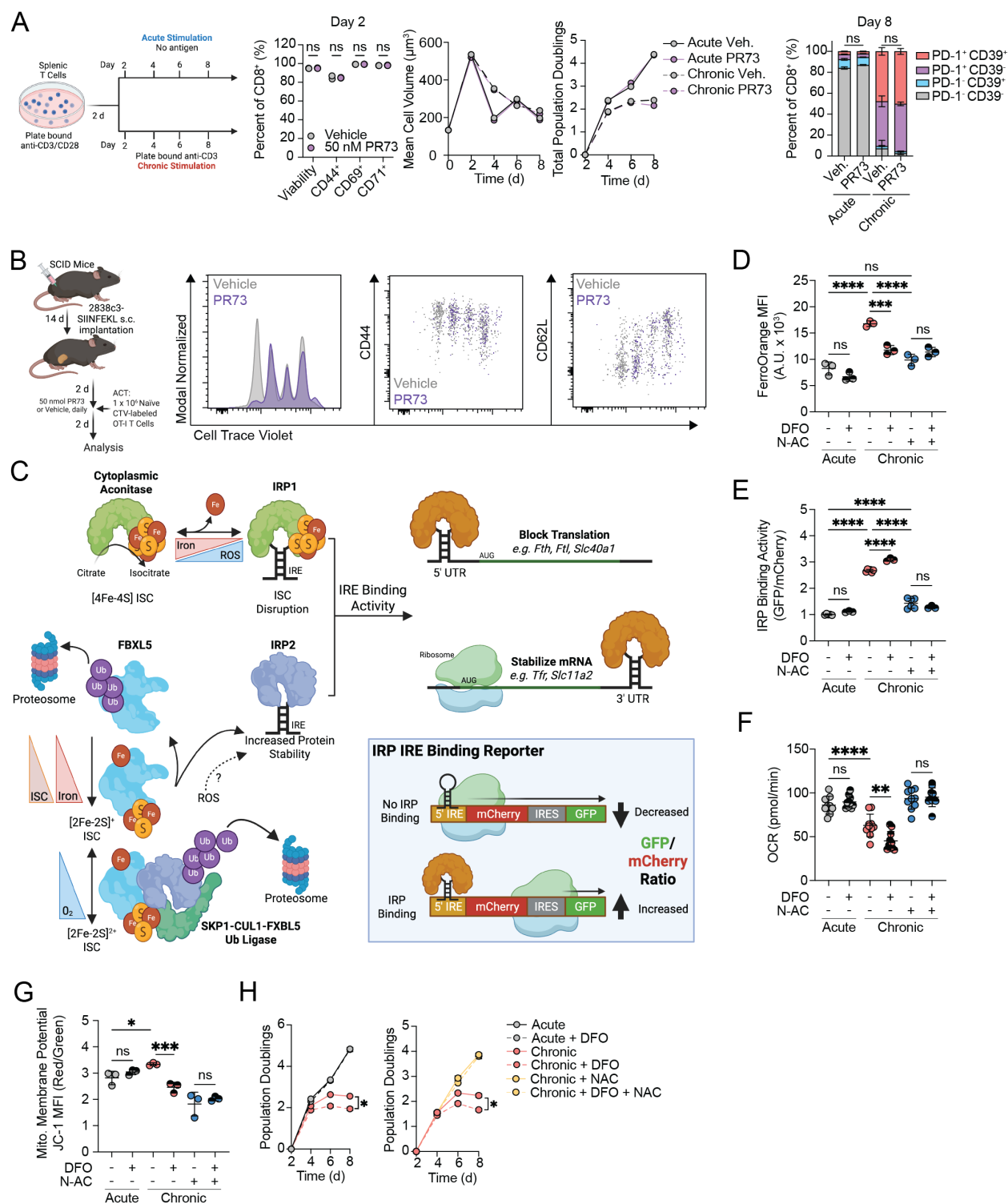

**Fig. S3. Related to Figure 3. Persistent antigen stimulation sensitizes CD8<sup>+</sup> T cells to iron restriction via a ROS-dependent mechanism. (A)** Platform of polyclonal splenic T cell activation *in vitro* culture under conditions of acute and chronic antigen stimulation. Immunophenotype, cell growth, and cumulative population doublings of CD8<sup>+</sup> T cells at specified timepoints when cultured in the presence or absence of PR73 (50 nM). **(B)** Following

establishment of 2838c3-SIINFEKL expressing tumors in syngeneic immunodeficient SCID mice, PR73 (50 nmol, I.P, daily) or vehicle control was administered for 2 days prior to the adoptive transfer of $1 \times 10^6$ naïve, cell trace violet (CTV)-labeled transgenic, antigen specific OT-I T cells. Mice were administered PR73 or vehicle control for an additional 2 days before tumor draining lymph nodes (tdLN) were isolated and phenotyped by flow cytometry. (C) Schematic of the regulation of IRP1 and IRP2 by iron and ISC availability, oxygen tension, and ROS (16-22) and the associated design of the IRP IRE binding activity reporter. (D-F) (D) Labile $\text{Fe}^{2+}$, (E) IRE binding reporter activity, (F) basal oxygen consumption rate (OCR), (G) mitochondrial membrane potential, and (H) cumulative population doublings of acute and chronically (10 nM N4) stimulated OT-I $\text{CD8}^+$ splenic T cells cultured in the presence or absence of DFO (500 nM) and/or N-Ac (10 mM) starting at day 2. All error presented as mean ± S.D., pair-wise analyses by Student's t-test, multiple comparisons by one-way ANOVA with Holm-Šidák correction, cumulative population doublings by two-way ANOVA with Geisser-Greenhouse correction (ns = not significant, \*$p < 0.05$, \*\*$p < 0.01$, \*\*\*$p < 0.001$, \*\*\*\*$p < 0.0001$).

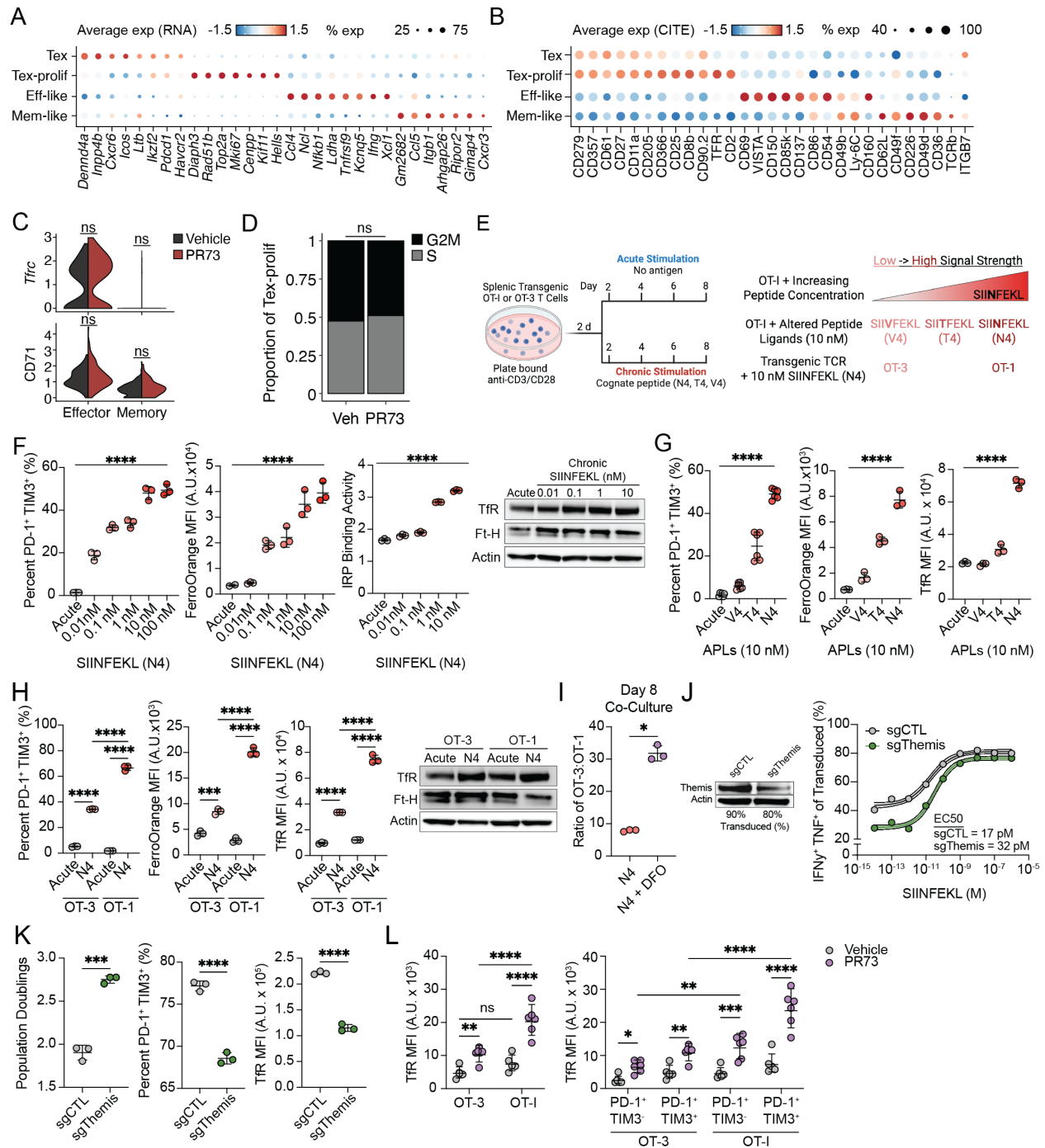

**Fig. S4. Related to Figure 4. TCR signaling strength calibrates disruptions in CD8<sup>+</sup> TIL iron metabolism.** (A-D) Adoptively transferred OT-I TILs from vehicle and PR73 treated (50 nmol, I.P., daily for 7 days) 2838c3-SIINFEKL tumor-bearing mice were isolated and analyzed using the DOGMA-seq workflow. Cluster defining (A) gene and (B) cell surface protein expression. (C) *Tfr* gene expression and Tfr surface protein expression in memory- and effector-like subsets stratified by treatment. (D) Transcriptomic cell cycle analysis of Tex-prolif TILs stratified by treatment. (E-H) (E) Pharmacological and genetic approaches to modulate TCR signal strength during persistent antigen stimulation *in vitro*. Cumulative population

doublings, labile $\text{Fe}^{2+}$, IRE binding activity, cell surface TfR expression, and representative western blots at day 8 of culture of (F) activated OT-I T cells cultured in the presence of increasing concentrations of SIINFEKL (N4) peptide, (G) OT-I T cells cultured with 10 nM APLs of varying avidity for the OT-I TCR (V4<T4<N4), and (H) OT-I and OT-3 T cells cultured in the presence or absence of 10 nM N4. For (F) and (G), p-value by one-way ANOVA with test for linear trend. (I) Following activation, OT-I and OT-3 T cells were co-cultured (1:1) for 6 days in the presence of 10 nM N4 with or without 500 nM DFO. Ratio of viable OT3:OT-I T cells at day 8. (J) Representative western blot of OT-I Cas9 T cells transduced with guide RNA targeting *Themis* (*sgThemis*) or control (*sgCTL*) guide at day 4 of culture. Dose response curves of acutely activated *sgCTL* and *sgThemis* OT-I Cas9 T restimulated at day 8 with increasing concentrations of N4 peptide (bands represent 95% CI). (K) Cumulative population doublings, labile $\text{Fe}^{2+}$, and surface TfR expression of *sgCTL* and *sgThemis* OT-I Cas9 T cells cultured for 6 days in the presence of 10 nM N4. (L) Surface TfR expression of OT-I and OT-3 TILs isolated from 2838c3-SIINFEKL tumor-bearing mice treated with vehicle or PR73. All error presented as mean ± S.D. Unless otherwise noted, pair-wise analyses by Student's t-test, multiple comparisons by one-way ANOVA with Holm-Sidak correction (ns = not significant, \*$p < 0.05$, \*\*$p < 0.01$, \*\*\*$p < 0.001$, \*\*\*\*$p < 0.0001$). Statistical analyses of DOGMAseq data are described in the Supplementary Methods.

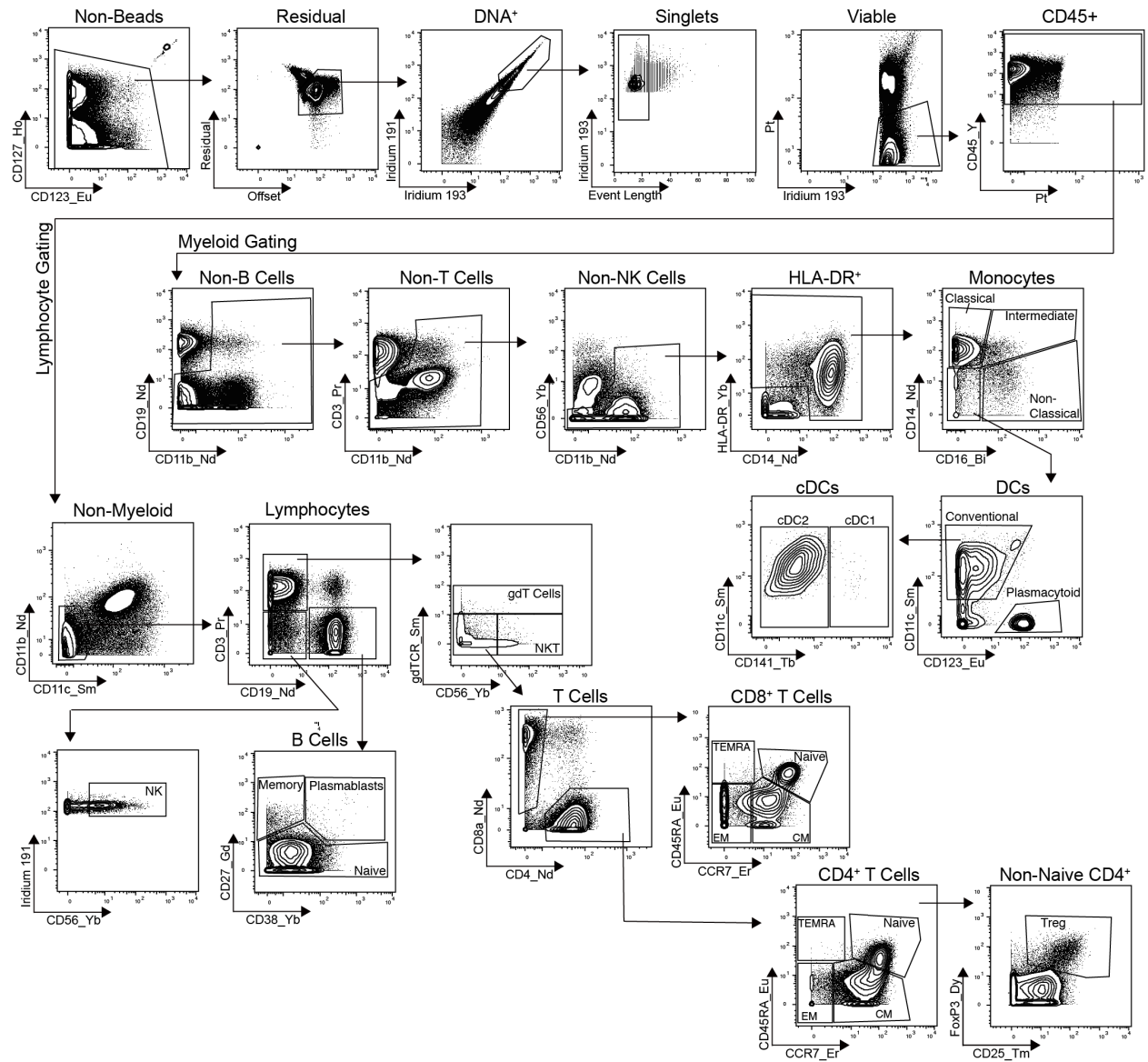

**Fig. S5. Representative CyTOF gating of PBMCs from patients on the PRINCE trial.**

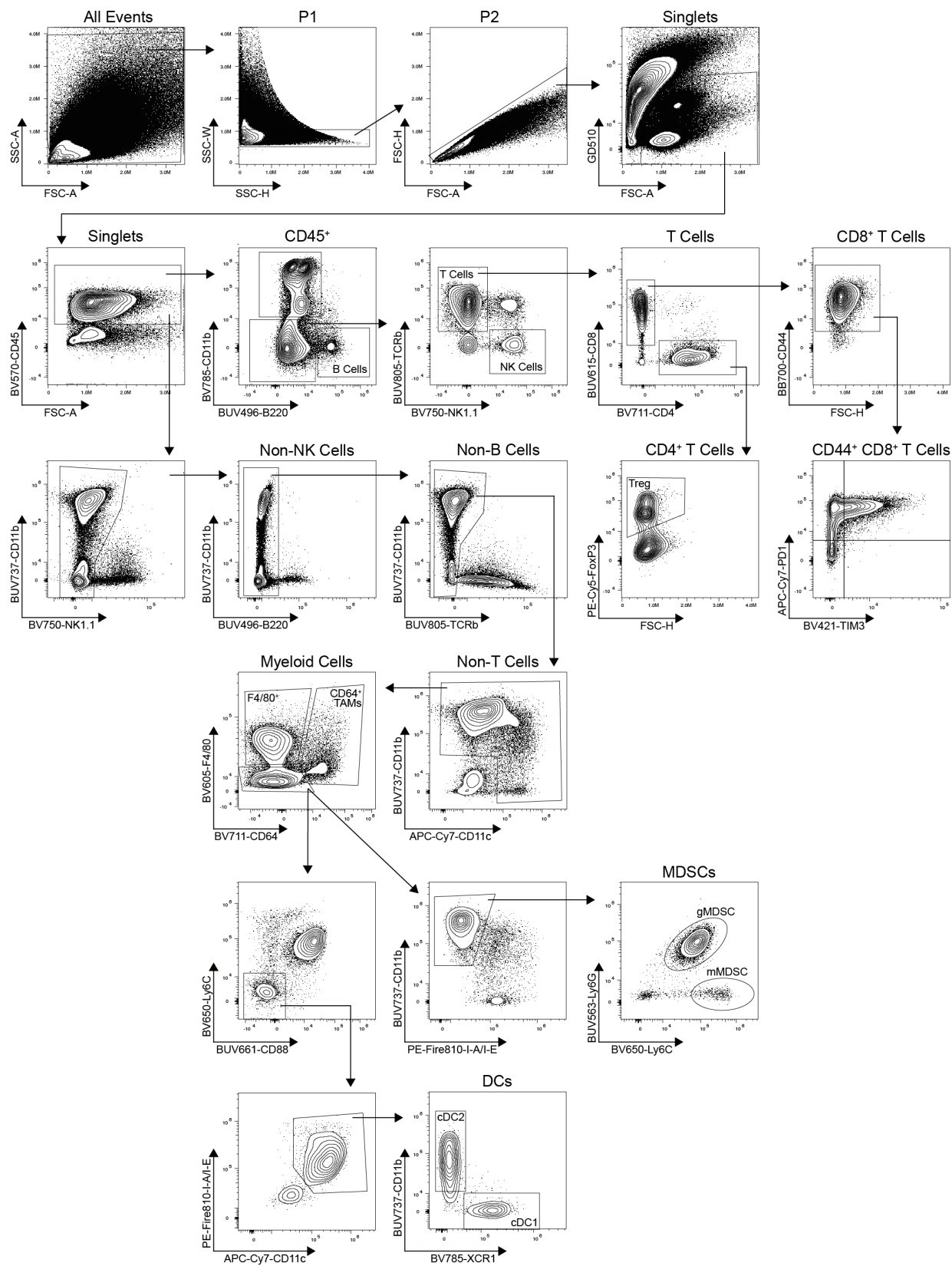

**Fig. S6. Representative flow cytometry gating strategy of murine tumors.**
